# CHEMOKINE SIGNALING IN BREAST CANCER: FOCUS ON CC-MOTIF CHEMOKINE LIGANDS

**DOI:** 10.1101/2024.03.01.583061

**Authors:** Sedat Kacar

**Affiliations:** Indiana University Melvin and Bren Simon Comprehensive Cancer Center, Vera Bradley Foundation Center for Breast Cancer Research, Department of Surgery, Division of Oncologic Surgery, Indiana University School of Medicine, Indianapolis, IN 46202, USA; Indiana Center for Regenerative Medicine & Engineering, Department of Surgery, Indiana University School of Medicine, Indianapolis, IN 46202, USA; Indiana University, Division of Pulmonary Medicine, Pulmonary, Critical Care and Sleep Medicine

**Keywords:** Breast Cancer, human CC-motif chemokines, CCR, chemokines, chemokine ligands/receptors, tumor initiation, immunogenicity

## Abstract

Based on 2020 data from GLOBOCAN, breast cancer holds the highest incidence rate in the majority of countries (159 out of 185 countries) and is the leading cause of mortality in 110 countries Chemokines are 8-10 kDa proteins activating through transmembrane G-coupled receptors (GPCRs) with 24 CC-motif type known human cytokines and 10 respective receptors. Chemokines and chemokines receptors have the potential for new breast cancer therapeutics modalities. In the complex chemokine world, not only one chemokine can bind to multiple receptors but also one receptor can be activated by several chemokine ligands. The fact that immune cells possess the potential to express multiple receptors and also secrete a diverse number of chemokines further intensifies the intricate and challenging to grasp chemokines interaction with receptors and microenvironment. In addition, the implication of chemokines in the malignant activation of tumor cells makes them indispensable contributors to the cancer microenvironment and influences the fate of tumor cells and the overall survival of the patients. Therefore, herein, we aimed in this study at attracting the researchers’ attention to the particular function of all known CC-motif chemokines and to-date findings, their expressional changes in normal, tumor and tumor adjacent tissues, important key effects in breast cancer and more importantly, we analyzed the breast tumor samples’ expression data retrieved from TCGA, Xena UCSC as well as UALCAN database regarding to showing any potential to be a general, subclass-specific or stage-specific marker as well as effects of overall survival of the patients and their correlation with literature findings. Basically, we desire this study to be a sort of reference insights for effects of human CC-motif chemokines in breast cancer or provide ready-to-use source and insights for researchers with the graphs and analyzes from different databases.

## 1. Introduction

Breast cancer is a leading female malignancy. Based on 2020 data from GLOBOCAN, the incidence of breast cancer in females keep the lead with a 3-fold difference to the second leading cancer type, colorectum cancer (Sung et al., 2021). Number of new cases in 2020 including all cancer types and both sexes, breast cancer settled in the leading position with 2,261,419 new cases, surpassing lung cancer (See **Fig. 1**). Of all cancer types, breast cancer has the highest mortality ratio in females, which is followed by lung cancer. When we consider both sexes, its rank is second after lung cancer according to age-standardized rate (ASR). Among women, 1 out of 4 cancer or 1 out of 6 cancer deaths stems from breast cancer. It holds the highest incidence rate in the majority of countries (159 out of 185 countries) and is the leading cause of mortality in 110 countries.

**Figure 1.**
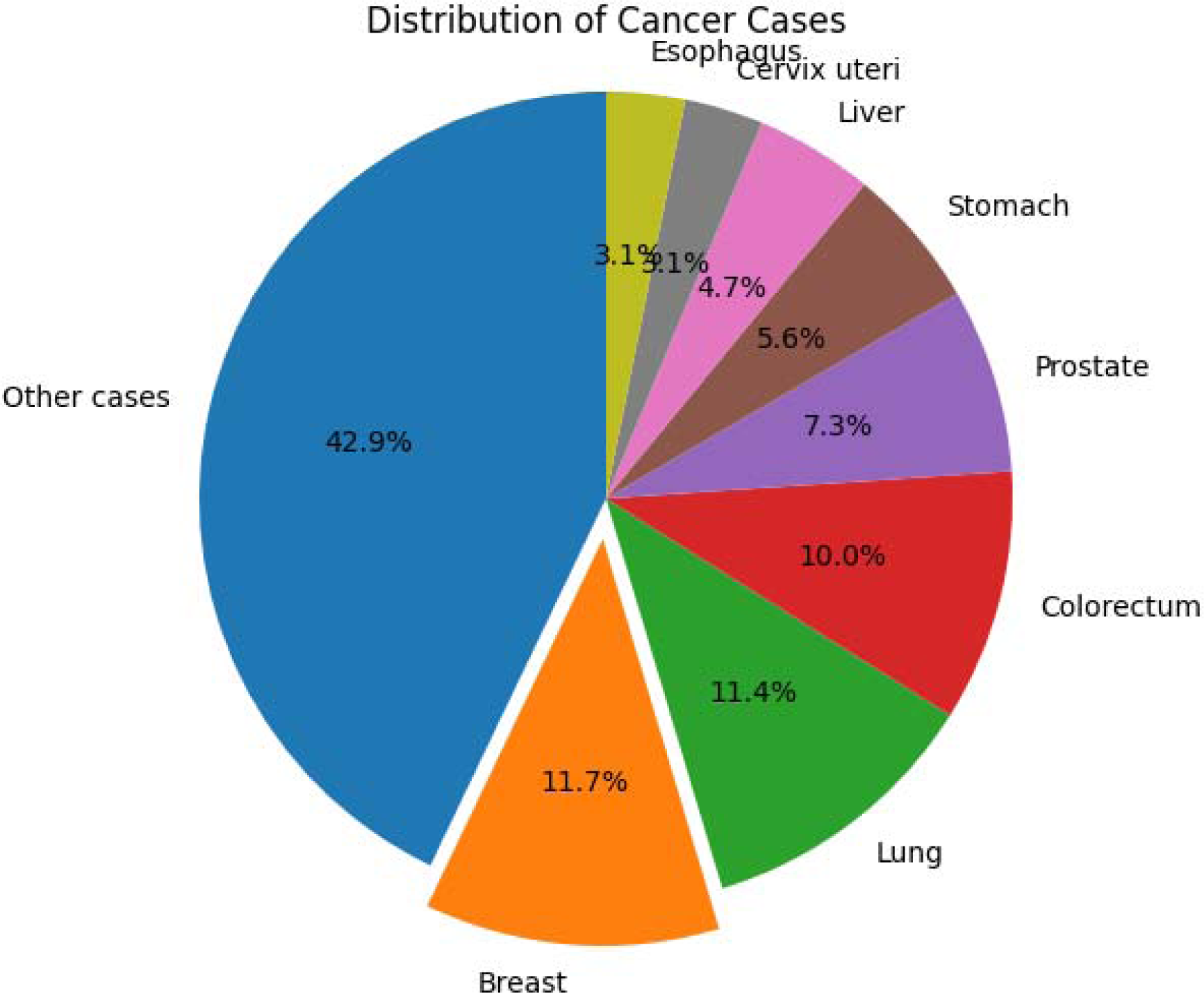
Number of new cases in 2020 from GLOBOCAN.

Chemokines are 8-10 kDa proteins that turn the target cells on through transmembrane G-coupled receptors (GPCRs). There are 30 (with subtypes) CC chemokine types. 4 of them including CCL6, CCL9/10, CCL12 and CCL21ser are only mouse-type chemokines. 8 of them including CCL3L1, CCL4L1, CCL13-16, CCL18 are only human-types CCLs. All other 18 CC-type CCLs are found both in mice and humans. To date, 10 different CC-motif chemokine receptors have been identified. The receptor location of 16 human CC-motif chemokines is on chromosome 21, with 2 on chromosome 22, 2 on chromosome 12, and 1 on chromosome 27.

Chemokines are basically tasked with the positioning and migration of immune cells during inflammation and infection. Other than that, in cancer, chemokines are notorious for being the cause of the bad prognosis of tumors by contributing to tumor angiogenesis and facilitating tumor cell migration, thereby metastasis. Intriguingly, in the chemokine world, many chemokines can bind to more than one receptor, and many chemokine receptors can activate various chemokine ligands, and that immune cells not only secretes various types of chemokines but also express multiple chemokine receptors, which all in all makes chemokine ligand-receptor interaction as focus-worthy as it is intricate and challenging. This promiscuous feature of the chemokine world becomes even more challenging to explore due to the fact that a chemokine may elicit diverse downstream signaling cascades depending on the triplet interaction among the target cell, chemokine subtype and the receptor it binds. In addition, the location, timing and duration of chemokine expression are other factors to be considered to effectively sort out the place of chemokines in cancer phenomenon, contributing to the complexity (Mollica Poeta, Massara, Capucetti, & Bonecchi, 2019) (See **Fig. 2B**)

**Figure 2.**
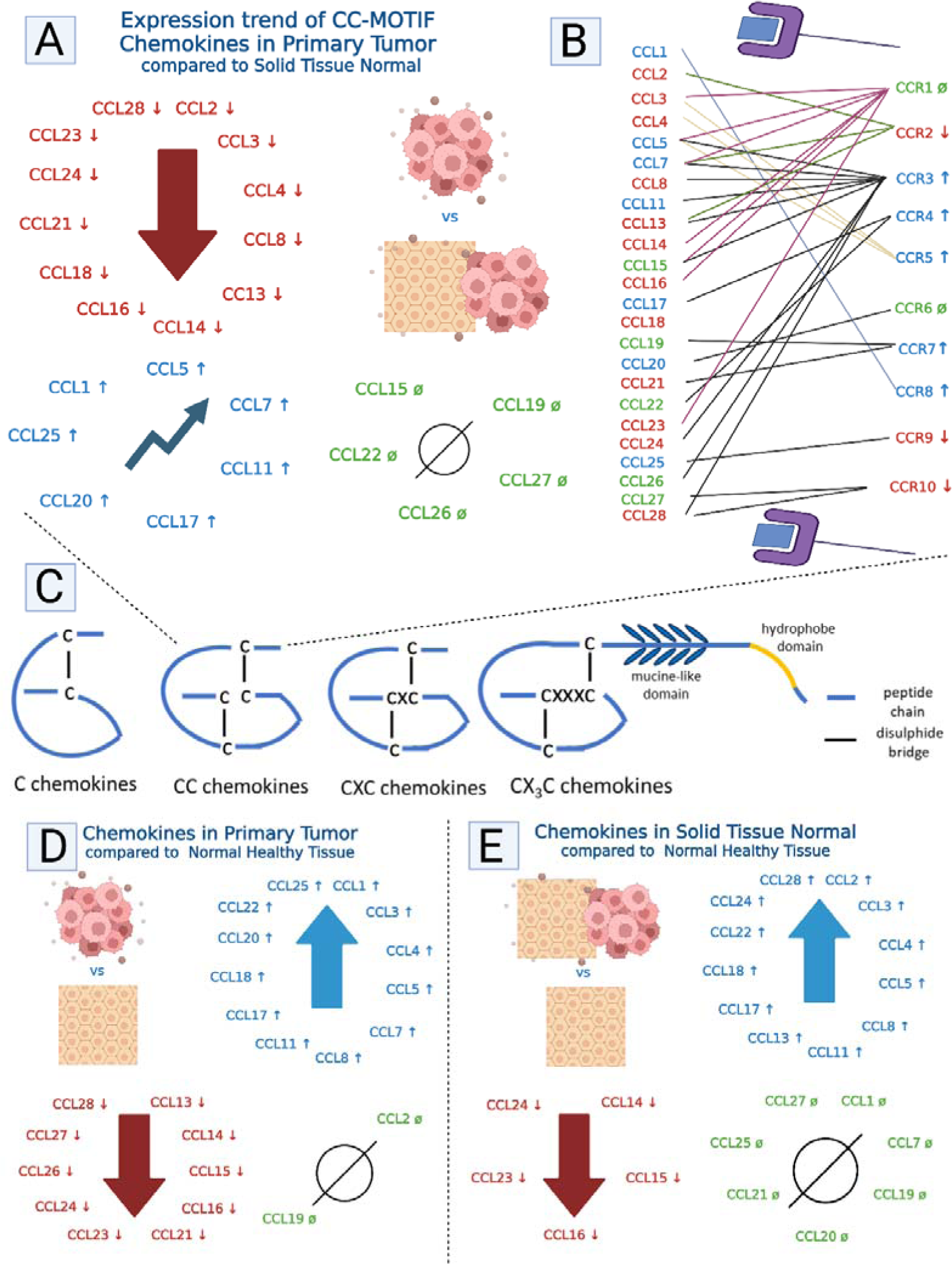
Expressional trends of all 24 human CC-type chemokines in tumor tissue when compared to normal solid breast tissue: Green(ø), blue (↑) and red (↓) denote “no change”, “a significant increase” and “ a significant decrease when compared to normal tissue adjacent to tumor (NAT) (**A**), corresponding CC-motif chemokine receptors to human CCLs: Green(ø), blue (↑) and red (↓) denote “no change”, “a significant increase” and “ a significant decrease when compared to NAT (**B**), structures of chemokines’ types (**C**), CC-motif chemokine human CCLs in solid tumor tissue: Green(ø), blue (↑) and red (↓) denote “no change”, “a significant increase” and “ a significant decrease when compared to healthy breast tissues (**D**) CC-motif chemokine human CCLs in NAT: Green(ø), blue (↑) and red (↓) denote “no change”, “a significant increase” and “ a significant decrease when compared to healthy breast tissues (**E**).

Chemokines and chemokines receptors have the potential for new breast cancer therapeutics modalities. Shedding light on this complex phenomenon will provide a wealth of benefits including insights about interactions and mechanistic trend change in tumor microenvironment. More importantly, this information will be harnessed to produce new and more robust treatments. Therefore, herein, we present this review to attract the researchers’ attention to the particular function of different CC-motif chemokines and to-date findings, important summarized tables as well as some critical topics. To be more focused and to have a point-to-target discussion, we only compiled the CC-motif chemokines, rather than including all cytokines and chemokines, ruling out any distraction.

According to the position of the first two N-terminal cysteine residues adjacent to the amino terminus, there are 4 types of chemokine families: C, CC, CXC and CX_3_C Chemokine Families (See **Fig 2C**). However, in this article we will be focusing on only CC-motif chemokines.

Basically, chemokines are a big family of small, secreted and structurally similar cytokines with chemotactic function, which takes a crucial role in the migration of the cells. Both cancer and stromal cells possess the cognate receptors of chemokines (Mollica Poeta et al., 2019). Chemokines both directly and indirectly modulate cancer cell growth by proliferating cancer cells and inhibiting their apoptosis. They are crucial for the host response to cancer by initiating a complicated migratory response that recruits leukocytes into the tumor milieu. They help the migration of tumor cells, which subsequently leads to metastasis. In addition, they have the tumor stromal cells release pro-tumorigenic factors like angiogenic and growth factors (Hughes & Nibbs, 2018). All in all, they do not just take a pivotal role in regular inflammation and immunity response but also serve as targets of oncogenic cascades and critical players in establishing the elements of tumor stroma.

Chemokines exert their effect through a chemokine gradient. It is a kind of concentration-dependent tumor cell attraction of chemokines. Epithelial secretions and signals constitute an environment that is full of chemoattractants with different gradients, forming an efficient platform for tumor cells to disseminate and expose to epithelial-mesenchymal transition (EMT). The interplay of chemokines with immunogenic cells could exhibit pro or anti-tumor impacts. Th1 immune response protects against cancer with antitumoral activity like NK cells, CD4+ Th1 and CD8+ cytotoxic lymphocytes. On the other hand, Treg and Th2 lymphocytes promote tumor proliferation (Mollica Poeta et al., 2019).

## 2. Material and Methods

### 2a. Retrieving database resources from TCGA

-TCGA-BRCA project was exploited from the NIH GDC Data Portal. Utilizing TCGABiolinks package program in R, TCGA data have been downloaded, prepared and gene expression data frame objects are constituted.CC motif chemokines are examined regarding their gene expression (FKPM). These results are reflected in Figures 3, 4 and supp. Fig 1. The packages were downloaded from “http://www.bioconductor.org/packages/release/data/annotation/html/org.Hs.eg.db. html” The difference in chemokine expression levels was exhibited by a column graph, which was drawn using R software (4.2.2). In addition, gene expression data from 999 samples were downloaded from TCGA database (https://cancergenome.nih.gov/); 904 tumor samples with no metastasis. and 88 normal solid tissue samples were used. Once all samples are downloaded, tumor and normal tissues are grouped. Briefly, samples with specific barcode codes with 999 samples are assigned with GDCquery function of TCGABiolinks. Then they were downloaded with GDCdownload function of the same package and prepared with GDCprepare fucnction with summarized Experiment argument. Thereafter, FPKM-based gene expression matrix is made. The names in the matrix, which was Ensembl based annotation are replaced with known gene symbols like CCL1,CCL2 etc. Then Normal and Tumor samples are assigned excluding metastatic samples. Base R function, boxplot is used to draw graphs with specific upper limits according to general trend of the data.

**Figure 3.**
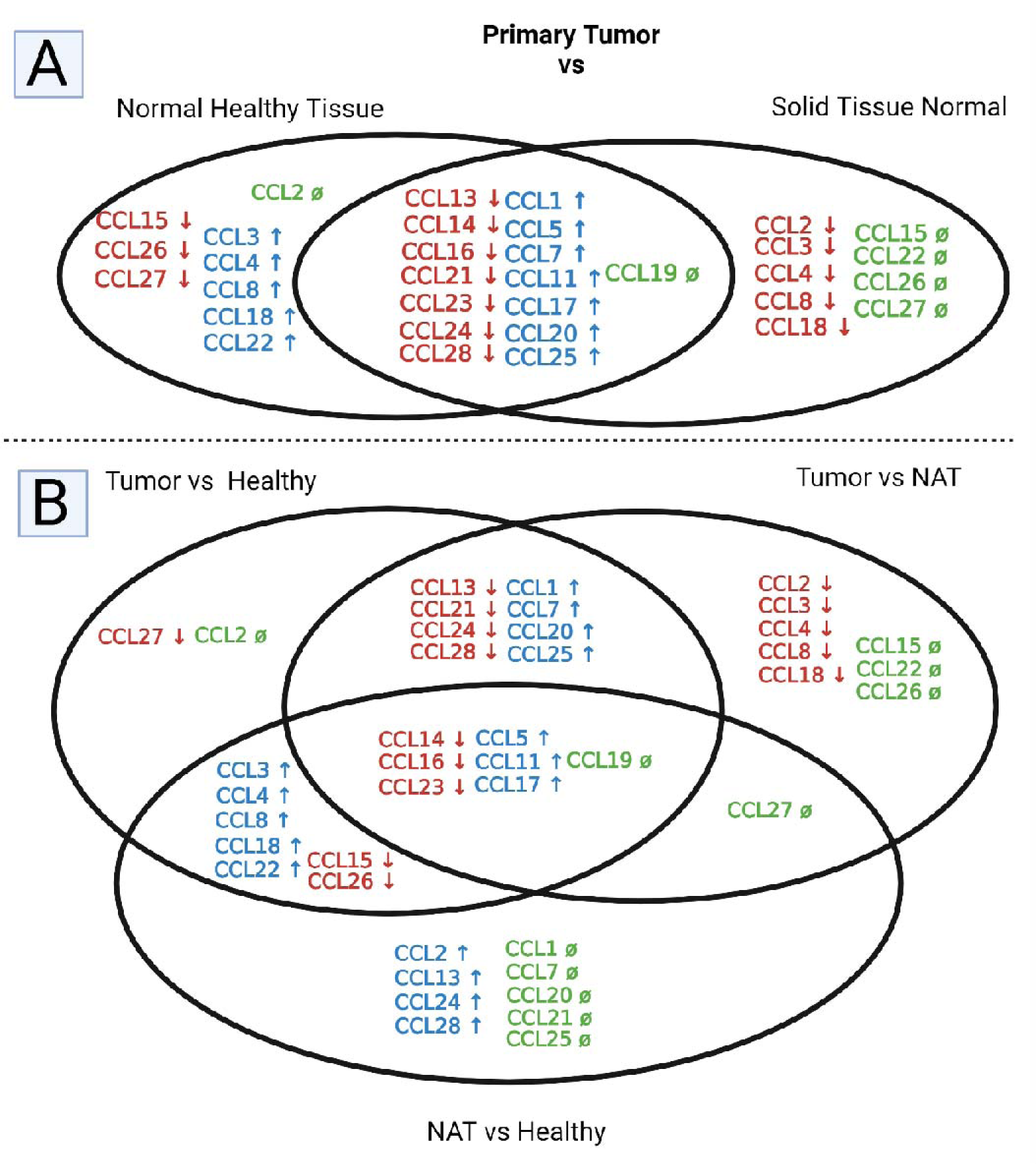
Venn Diagrams of human CC-motif chemokines: Comparing Expression Changes in Primary Tumor, Healthy Breast Tissue, and Normal Tumor Adjacent Tissue (NAT) (A) The Dual Venn Diagram of the overlapping similarities and distinctions in primary tumor tissue CCL expression changes over healthy breast tissue (left side) or NAT (right side). Green(ø), blue (↑) and red (↓) denote “no change”, “a significant increase” and “a significant decrease when compared to healthy (left side) or NAT (right side). (B) The triplet Venn Diagram of the overlapping similarities and distinctions in primary tumor tissue and NAT. In this diagram, primary tumor expression changes when compared to normal breast tissue (Tumor vs healthy) or tumor adjacent normal tissue (Tumor vs NAT) and NAT expression changes when compared to normal breast tissue (NAT vs Healthy) are shown. Green(ø), blue (↑) and red (↓) denote “no change”, “a significant increase” and “a significant decrease”, respectively.

**Figure 4.**
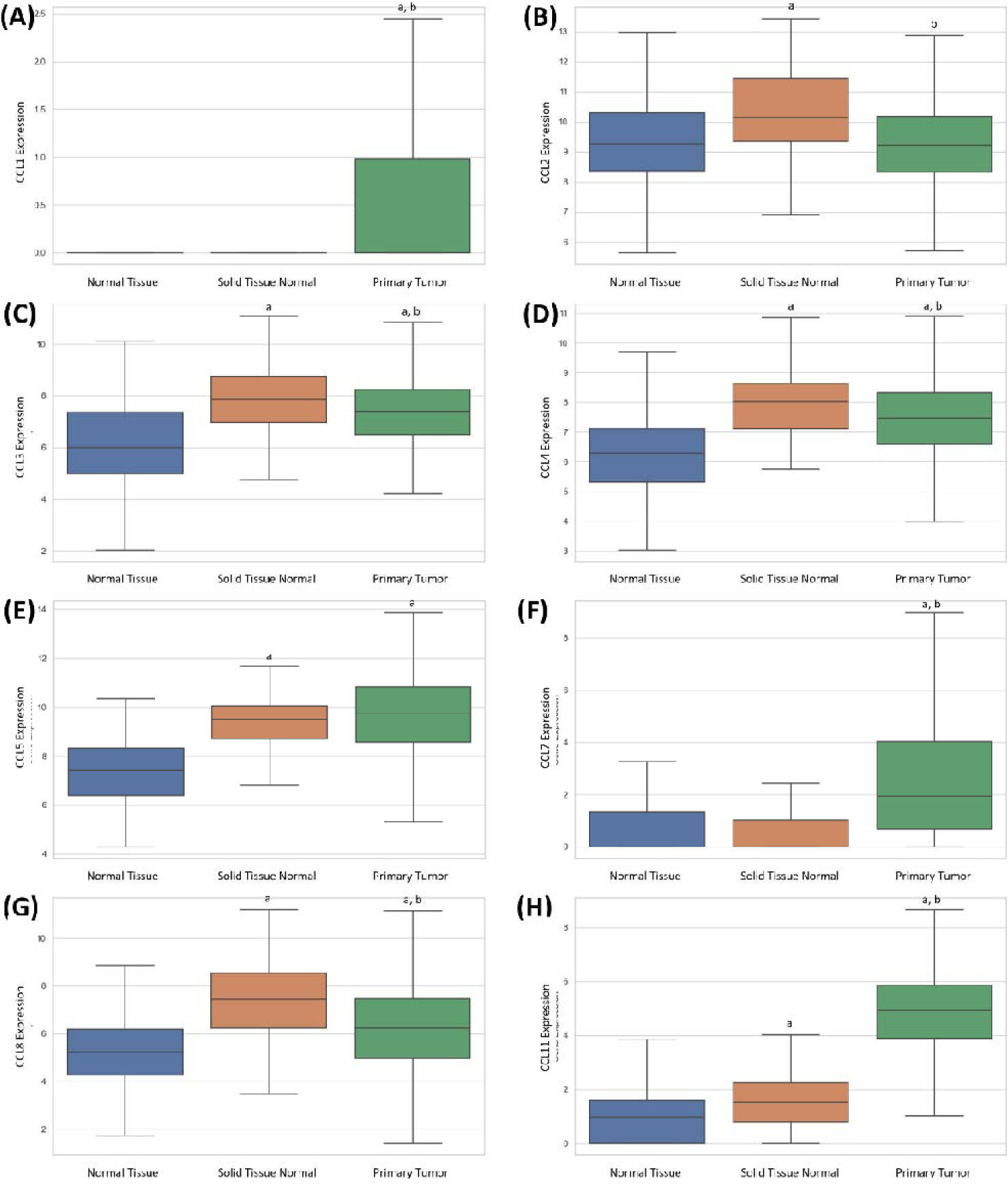
The expressions of CCLs 1-5, 8, 11 retrieved from Xena website and modified (n=178 for healthy normal tissue, n=113 for tumor-adjacent normal tissue, n=1092 for primary tumor). The unit of the expression values are log_2_(RSEM norm_count +1). **Note:** In all graphs, “a” and “b” symbols designate a significant difference of p<0.5 when compared to normal tissue and solid tissue normal, respectively based off of Tukey’s multiple comparison post hoc test.

### 2b. Using the ready-to-use graphs in UALCAN website

CCL1-5, 7-8, 11, 13-28 has been searched in TCGA motor of UALCAN website with picking up “Breast invasive Carcinoma” in the drop-down menu of TCGA dataset. In coming page, sample types, major subclasses and individual cancer stages in drop-down menu are downloaded for each CC-motif chemokines. Furthermore, in survival data was downloaded through survival data tabbed panel for each CC motif chemokines. Once the graphs have been downloaded from website one by one. The downloaded figures are merged and organized by using matplotlib.pyplot and matplotlib.image modules of matplotlib lilbrary of python 3.11.2 on Integrated Development Environment (IDE) of VS Code, which are represented in fig 5-9 and supp Fig 2 and 3. Ultimately, although all figures were ready to use, the labels on figures are improved and modified in PowerPoint. Statistical values on UALCAN website are used as they are.

**Figure 5.**
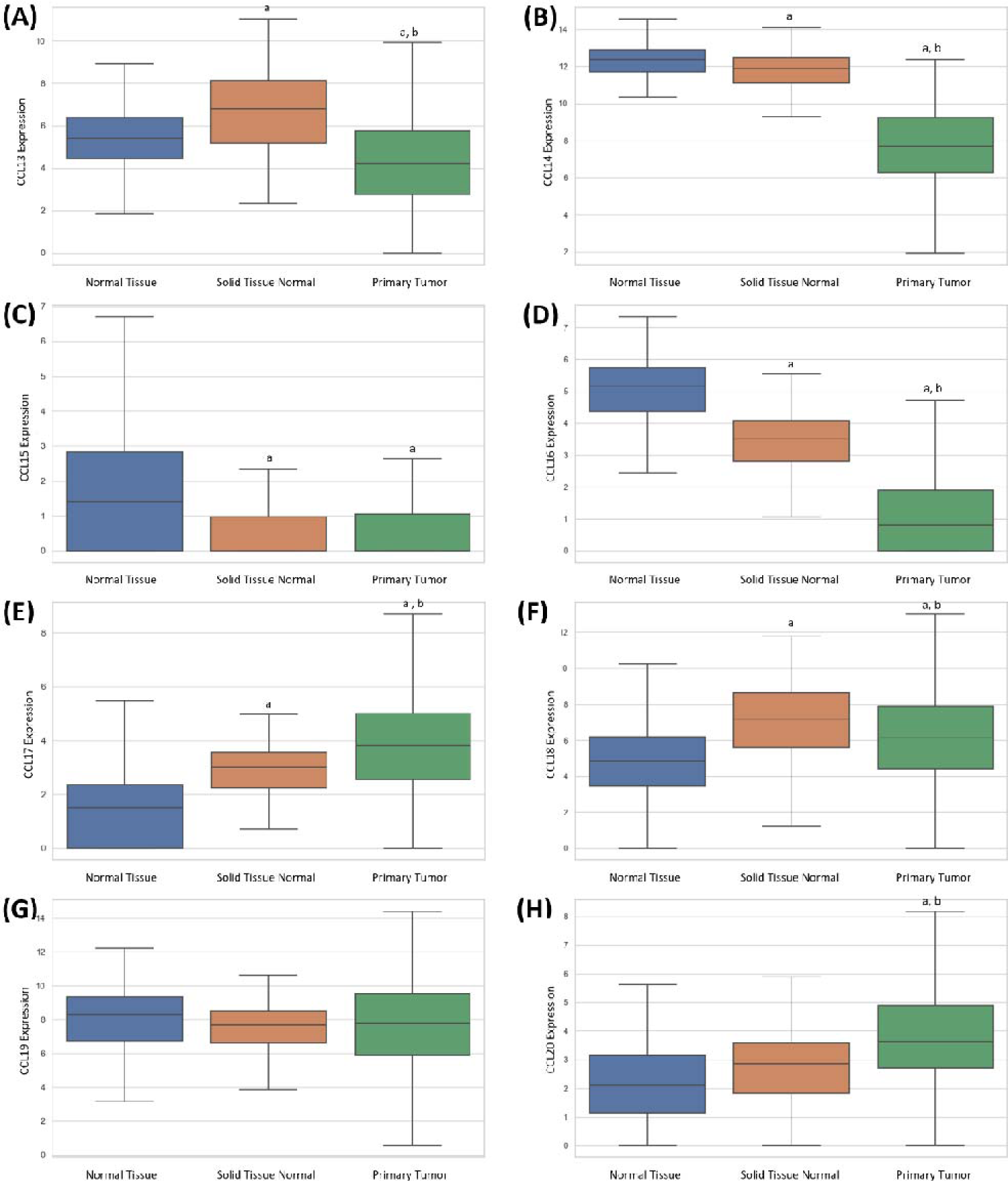
The expressions of CCLs 13-20 retrieved from Xena website and modified (n=178 for healthy normal tissue, n=113 for tumor-adjacent normal tissue, n=1092 for primary tumor). The unit of the expression values are log_2_(RSEM norm_count +1) **Note:** In all graphs, “a” and “b” symbols designate a significant difference of p<0.5 when compared to normal tissue and solid tissue normal, respectively based off of Tukey’s multiple comparison post hoc test.

### 2b. UCSC Xena website utilization for healthy normal breast cancer tissues

Since TCGA database only has tumor-adjacent normal tissue and that GTEX portals normal breast cancer tissue expression values are from different experimental sample processing than TCGA, we downloaded all human CCLs gene expression data is from https://xena.ucsc.edu/ as a tsv. files, where TCGA and GTEX expression data are re-aligned to hg38 genome and analyzed by RSEM method (Goldman et al., 2020). Thus, we reduced the possible batch effects minimum between different expression data. Data in tsv files are processed through Python 3.11.2 by using the packages of pandas, seaborn, matplotlib.pyplot, scipy.stats and statsmodels.stats.multicomp. Briefly, the tsv files are read, and the column information of the group names of normal breast, primary tumor and tumor adjacent normal tissues and their corresponding RSEM expression values are extracted. Then, the statistical test are applied by scipy.stats and statsmodels.stats.multicomp. Eventually, the graphs are generated and the graphs are modified by adding respective statistical results and re-writing the less-visible legends. The git.hub link of the sources codes are accesible in https://github.com/sedatkacar56/xenadataprepare.md.git

### 2c. Statistical analysis

The statistical analysis for the data retrieved from TCGA is performed accordingly. First the trend of group data and values were checked with Shapiro-Wilk test if it follows a normal distribution or not. For normally distributed data, dual statistical tests were done by t-test; otherwise, Mann Whitney test has been applied. Most of the gene’s distributions fell in the latter category and the Mann Whitney test was applied in TCGA retrieved data. The data retrieved from UALCAN already had its statistical data. For the data from UCSC Xena website, ANOVA multiple pairwise comparison test was utilized for all samples. p< 0.05 is deemed as statistically significant in all statistical analyzes.

### 2d.Literature review

Other than that, the CC-motif chemokines are scanned in literature and background information is shared in the article.

### 2. Reported Effects and Gene Expressions from TCGA of Different Human CCLs

Only Human CCLs are mentioned. Since CCL 6, 9, 10 and 12 are mouse-type CCLs, they have not been mentioned in the following discussion of this section. Gene expression data of TCGA and figures were gained from UALCAN website of https://ualcan.path.uab.edu/analysis.html (Chandrashekar et al., 2017; Chandrashekar et al., 2022). In the **Fig 2-3 and Supp. Fig 7**, the gene expression changes (upregulation/downregulation/no change) are summarized for all CC-motif huma chemokines as well as their respective receptors (CCR1-10). Summarized results and expression change trend of all CCLs are given in Fig. 2 as a summarized form. These charts can be utilized to see the clear picture of transcriptional trend in the triplet of healthy, tumor and tumor adjacent normal tissues.

In the TCGA database, the normal samples employed are tumor adjacent normal samples; therefore, we also downloaded the gene expression data of CCLs (as well as CCRs) from UCSC Xena website (https://xena.ucsc.edu/compare-tissue/). And analyzed this data with Python 3.11.2. What makes this website special is that they include a GTEX (Genotype-tissue Expression) portal database samples which are re-aligned, normalized and re-analyzed together with TCGA primary tumor and tumor adjacent tissue expression counts, making the analysis more robust and reliable by decreasing the batch effects. In our analysis, we made use of variuos databases and also manually retrieved and analyzed the data from GTCA utilizing TCGAbiolinks package in R. This approach enables the researchers to access all data and interpret results based on their discretion, minimizing the analyzer or website-based issues to the least. And we think the graphs, diagrams and schemes will offer the researchers who are exploring the effects of CC-motif chemokines on breast cancer an accessible m point.

## Immune Cells In Cancer

Before delving into the impact of CCL1 on cancer, let’s establish a foundation by understanding some key terms. Thus, coming texts would be clear, for example, when we say ’CCL1 increases regulatory T cells,’ it implies CCL1 has a role in promoting the activity of regulatory T cells, which are are known to inhibit cancer.

Prior to starting, it is essential to provide contextual information about T cell and immune cell types, elucidating in which way they affect the breast cancer. This preliminary insight will facilitate the interpretation of the the effects of CC-motif chemokines. For instance, the understanding that regulatory T cells have the potential to suppress cancer growth, CCLs recruiting these cells may have inhibitory effects or vice versa.

T lymphocytes are adept roamers of immune surveillance navigating and scrutinizing throughout the body for any foreign substances. With certain signals, they differentiate in such a way that they become distinct and mutually exclusive. T cells with TCR affinity for MHC-1 are CD8+ T cells and T cells with TCR affinity for MHC-2 are CD4+ T cells. Based on stimulatory cascade signals from cytokines and stromal cells, they could further specialize into T-helper or T-regulatory cells.(Raskov, Orhan, Christensen, & Gögenur, 2021).

### Tumor-infiltrating lymphocytes (TIL)

#### CD8+ cytotoxic T lymphocytes (CTLs)

They have TCR affinity for MHC1. Cytotoxic CD8+ T cells constitute the core of cancer immunotherapy, which are the powerful effectors in the anticancer immunity (Raskov et al., 2021). Therefore, we can consider them “beneficial contributors” within the context of cancer.

#### CD4+ T helper (Th) cells

They have TCR affinity for MHC2. According to cytokine spectrums and regulatory functions, classic CD4+ Th subsets are separated into 5 categorizations: Th1, Th2, Th9, Th17 and follicular helper T cells. CD4+ T cells were illustrated to contribute to the antitumor immunity and directly eliminate tumor cells through cytolytic mechanisms. (Li, Wu, Yang, Zhang, & Jin, 2020). Therefore, we can consider them “beneficial contributors” within the context of cancer.

### Regulatory T cells (T_regs_)

Regulatory T (Treg) cells serve a critical purpose in immune homeostasis by suppressing excessive immune responses. The common molecular marker of those cells is FOXP3, which is a key regulator gene of Treg cells. In the realm of tumor immunity, Treg cells, recruited to the site by certain chemokines such as CCLs 1, 17/22, 28, 9-11 hinder the body’s defense to tumors, assisting tumor growth. Regulatory T cells are then activated and inhibit antitumor immune responses. A conspicuous influx of Treg cells is correlated with poor survival across diverse types of cancer. (Kos & de Visser, 2021). In summary, Tregs are considered “detrimental contributors” within the context of cancer.

### Tumor-associated macrophages (TAM) and Cancer-associated fibroblasts (CAF)

TAMs induce angiogenesis and suppress immunity by secreting VEGF-A and upregulating the production of TGF- and IL-10 and prevent CD8+ cell infiltration. Likewise, CAFs evoke angiogenesis and generate a dense tumoral micro-environment. Tumor microenvironment with abundant TAMs and CAFs is associated with poor immune response as it is hostile to lymphocyte recruitment (Monteran & Erez, 2019; Raskov et al., 2021). Thus, both TAMs and CAFs are considered “detrimental contributors” within the context of cancer.

## C-C motif chemokine ligand 1 (CCL1)

CCL1 is also known as I-309, and its only known receptor is CCR8 (Fig 2B). One of 7 increased chemokines in solid tumor compared to normal based on TCGA data analysis (Fig. 2). In one study where 199 invasive breast cancer tissues were stained immunohistochemically, CCL1 is shown to be expressed in the majority of breast cancer tissues. Intramural CCL1 expression was observed in higher-grade tumors (G4). CCL1 was proposed to have a correlation with regulatory T cells based on the results of FoxP3+ staining (Kuehnemuth et al., 2018). CCL1 expression fosters the activation of regulatory T cells, which accelerates tumor development, concomitantly hampering cytotoxic T cells to initiate an anti-cancer immune response.

As shown in Figure 2, 3, 4 and Supp. Fig 1, CCL1 expression of primary solid tumor is significantly higher than healthy breast tissue and NAT samples. In addition, there is no significant difference between tumor normal and healthy normal tissue samples. It seems solid tumor shows a differential expression of CCL1, which is consistent with literature findings.

In Fig 5A, although no eye-catching difference is visible on the graph, a significant difference between tumor and NAT samples are recorded. This difference is more visible and conspicuous in TNBC subtype (Fig 5B), which is a more invasive type of breast cancer. In early stages of cancer, namely stage 1 and 2, higher CCL1 levels are observed. On the other hand, the CCL1 levels of stage 4 are similar to NAT (Fig 7C). In Kaplan Meier survival plot, survival plot of the patients with high CCL1 expression indicates a longer survival plot up to 17 years than low expressed groups (p<0.01). This survival plot of CCL1 is the one showing the strongest significant value among other human CC-motif chemokines (Supp. Fig 5A).

**Figure 6.**
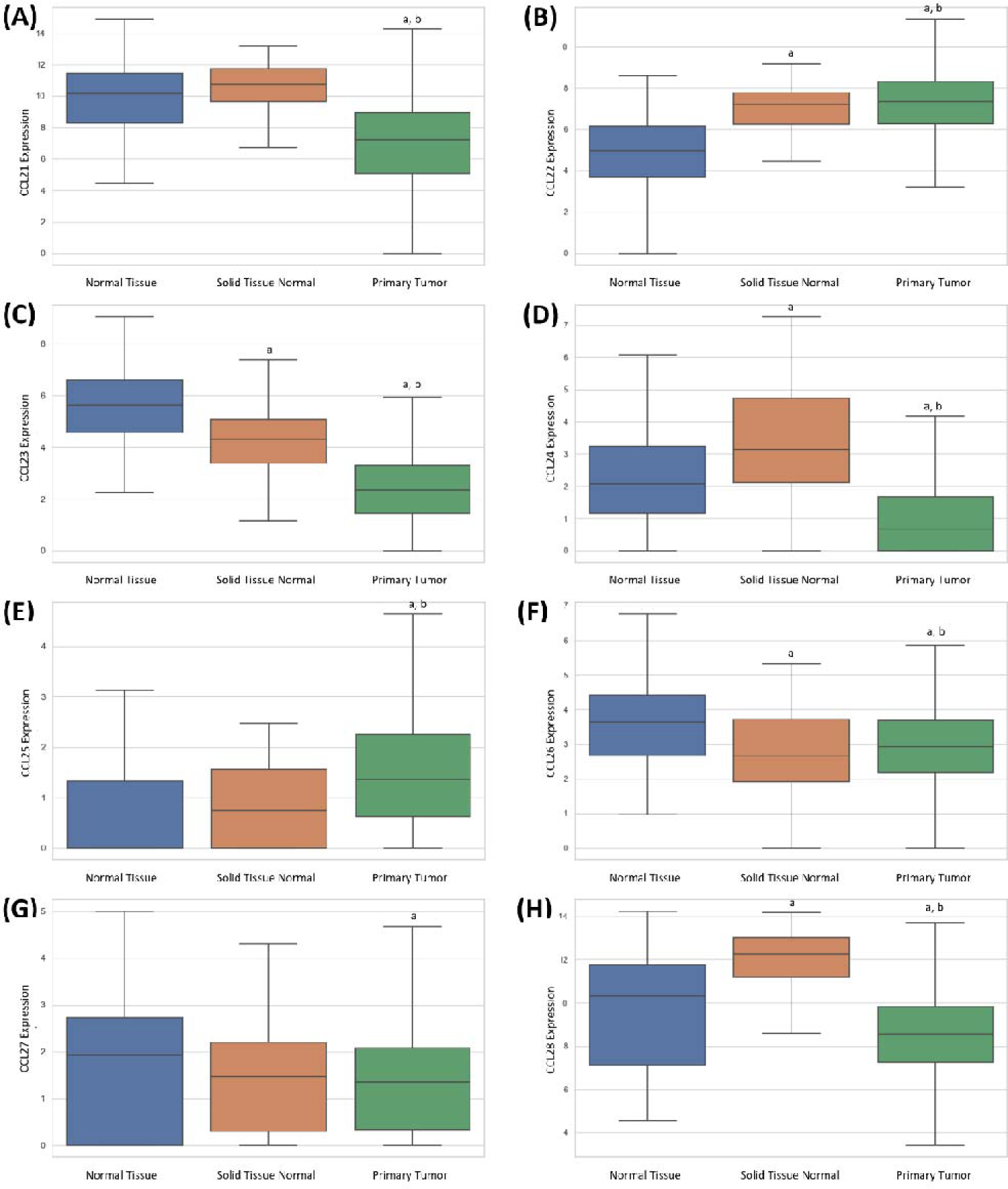
The expressions of CCLs 20-28 retrieved from Xena website and modified (n=178 for healthy normal tissue, n=113 for tumor-adjacent normal tissue, n=1092 for primary tumor). The unit of the expression values are log_2_(RSEM norm_count +1). **Note:** In all graphs, “a” and “b” symbols designate a significant difference of p<0.5 when compared to normal tissue and solid tissue normal, respectively based off of Tukey’s multiple comparison post hoc test.

**Figure 7.**
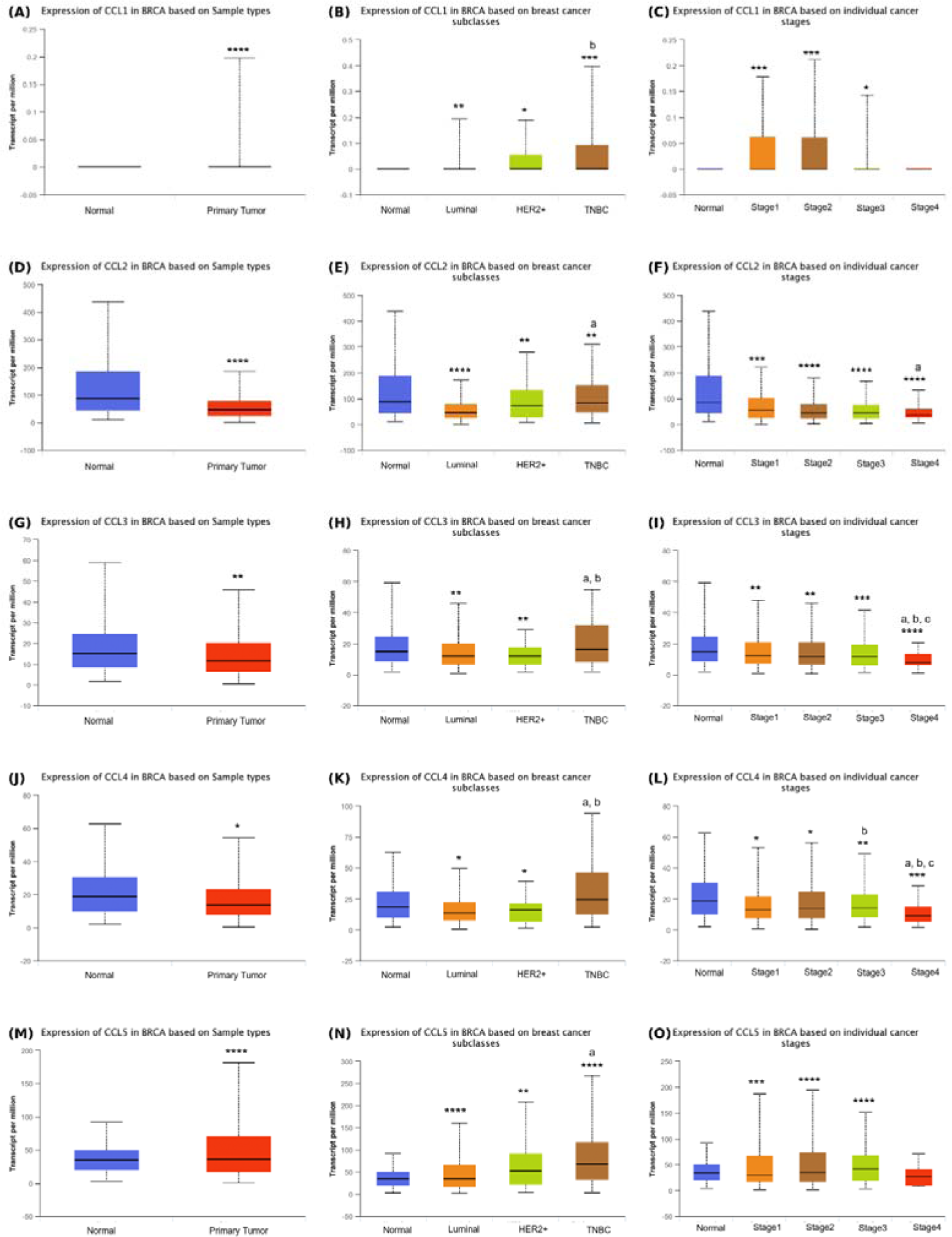
The expressions of CCLs 1-5 from TCGA database. *Sample type*: The expressions of CCL1 (**A**), CCL2 (**D**), CCL3 (**G**), CCL4 (**J**) and CCL5 (**M**) in normal (n=114) and breast cancer tissue samples (n=1097). *Breast cancer molecular subclasses*: The expressions of CCL1 (**B**), CCL2 (**E**), CCL3 (**H**), CCL4 (**K**) and CCL5 (**N**) in normal tissue (n=114) and breast cancer subtypes of luminal (n=566), HER2^+^ (n=37) and triple negative (TNBC, n=117). *Cancer stages:* The expressions of CCL1 (**C**), CCL2 (**F**), CCL3 (**I**), CCL4 (**L**) and CCL5 (**O**) in normal tissue and different stages including stage 1 (n=183), stage 2 (615), stage 3(n=247) and stage 4 (n=20). **Note:** In all graphs, *, **, *** and **** symbols designate a significant difference of p<0.5, p<0.01, p<0.001 and p<0.0001, respectively. In the individual cancer stages graph, the small letters of a, b and c designate a significant difference between stages 1, 2 and 3, respectively. In the breast cancer subclasses graph, the small letters of a and b designate a significant difference between luminal and HER-positive subclasses, respectively.

## C-C motif chemokine ligand 2 (CCL2)

CCL2 is also known as MCP-1, and its main known receptor is CCR2, but it was also reported to bind CCR4, whereby provoking breast cancer development (Xu, Wang, Xia, Wei, & Wei, 2021)(Fig 2B). However, CCR4 was also reported to not display receptor activities for CCL2, CCL3, or CCL5, as a results of binding, chemotactic, or calcium mobilization assays (Yoshie, 2021). CCL2 is documented to play a role in malignant behaviour of breast cells (Soria & Ben-Baruch, 2008). Fibroblast-secreted CCL2 boosted the filtration of innate immune cells and neo-vasculature. (PMC3778011). As a marker known as TIL (tumor-infiltrating lymphocyte), CCL2 is found to be more prevalent in ER-negative cancers than LumA subtype cancers (Franzen et al., 2019).

As shown in Figure 2, 3, 4 and Supp. Fig 1, CCL2 expression shows different differential expressions in Tumor/Healthy, Tumor/NAT and NAT/Healthy tissues comparisons. While it does not show any difference between tumorigenic and healthy breast tissue samples, it shows higher expression values in NAT tissue. It is conceivable that CCL2 might have a crucial function in the tumor microenvironment. Its heightened expression in the tumor-adjacent normal tissue might imply the presence of the nearby tumor. However, the lack of significant difference between tumor and healthy breast tissues regarding this chemokine is obscure and unknown. According to our analysis, that is the only CC-motif chemokine in human that shows a significant increase in NAT with no difference between tumor and healthy tissue.

According to TCGA data (Fig 7 D-F and Supp. Fig 2B), there is a highly significant difference between tumor and NAT samples (Fig 7D, p<0.0001), as we mentioned above. This significant difference between NAT samples persists in each subgroup in cancer subclasses and in different stages of the cancer. What’s more, an ascending trend in CCL2 is observed toward more invasive cancer subtype (from luminal subtype to TNBC). CCL2 levels of TNBC are lower than NAT samples but higher than luminal subtype of breast cancer. In cancer stages, a general decrease in CCL2 from stage 1 to 4 is evident (Fig 7D-F). There are not any significant differences or any visible trend in overall survival curves between the samples with high vs low expression of CCL2 (Supp. Fig 5B).

## C-C motif chemokine ligand 3 (CCL3)

Chemokine ligand 1 (CCL3) is also known as MIP-1α, and its known receptors are CCR1 and CCR5. Docetaxel (DTX) is an antimitotic chemotherapy agent, broadly utilized in prostate, lung as well as breast cancers. CCL3 chemo-sensitizes breast cancer cells to docetaxel by activating M1 macrophages and enables phagocytosis of cancer cells. High CCL3 is associated with better DTX chemosensitivity and better prognosis (Sheng et al., 2022).

As shown in Figure 2, 3, 4 and Supp. Fig 1, CCL3 expression is significantly higher both in tumor and NAT than healthy breast tissue. When compared to tumor and adjacent normal tissues, it is clearly seen that NAT heightened CCL3 expression. The increased picture of CCL3 expression in NAT might imply the more crucial role of CCL3 in tumor microenvironment. It can be employed as a clear-cut marker for the diagnosis of breast cancer. Let’s also check its trend in terms of cancer subtype and stages below.

According to TCGA data (Fig 5 G-I and Supp. Fig 2C), primary tumor tissue shows significantly lower levels of CCL3 than those in normal breast tissue (Fig 5G). While luminal and HER2+ breast cancer subtypes exhibit a significant diminished levels of CCL4 (p<0.01) relative to normal samples, TNBC shows no difference. However, CCL4 expression levels of TNBC subtype were significantly higher than both luminal and HER2+ subtypes (Fig H). Regarding cancer stages, a smooth decreasing trend is visible from initial stages to late stages in CCL3 levels. The cancer stage 4 differed significantly from all 3 stages in addition to normal tissue samples (Fig 3I). In Kaplan Meier Survival plot, no significant difference between high and low CCL8-expressing patients is recorded. (Supp. Fig 5C).

## C-C motif chemokine ligand 4 (CCL4)

Other names of CCL4 are macrophage inflammatory protein-1β and MIP-1β. It behaves as a chemoattractant of various immune cells and its receptor is CCR5, expressed on immature DC and TH1-polarized T cells (Nguyen-Hoai et al., 2016). With its cognate ligand, CCL4 is suggested to induce tumor growth, included as one part of the CCR5-dependent chemokine axis and considered a potential noteworthy therapeutic target in advanced-stage breast cancer. In the original study, according to the clinical data of healthy control-stage II and stage III patients shared, CCL4 seems to increase in stage II and then retrogress again to normal control levels in stage III (Pervaiz et al., 2019).

As shown in Figure 2, 3, 4 and Supp. Fig 1, CCL4 expression increases prominently both in tumor and NAT relative to healthy breast tissue samples. When compared to tumor and adjacent normal tissues, we also see that NAT has a significantly higher value of CCL4 expression. The pronounced presence of CCL4 in NAT points towards an enhanced role within the context of the tumor microenvironment. It has the potential to be utilized as a marker for diagnosing breast cancer.

According to TCGA data (Fig 5 J-L and Supp. Fig 2D), CCL4 levels are less in primary tumor breast tissue than in normal breast tissue. Among molecular subtypes, luminal and HER2+ breast cancer subtypes indicate a significantly low CCL4 levels (p<0.05) when compared to normal tissue. On the other hand, TNBC subtype did not show any difference from normal tissue. However, CCL4 expression levels of TNBC subtype were significantly higher than both luminal and HER2+ subtypes, implying CCL4 to be a future subtype related marker. Comparing each cancer stage with normal breast tissue samples, it is plausible that CCL4 expression is significantly lower in each cancer stage and there is a decreasing trend from early stages toward late stages. The latter has the significantly lowest levels of CCL4 when compared to all other stages (Fig 5L). In Kaplan Meier Survival plot, no relation between high and low CCL4-expressing patients is apparent at all. Both curves virtually follow an overlapping line (Supp. Fig 5D).

## C-C motif chemokine ligand 5 (CCL5)

This C-C motif ligand is also known as RANTES and its receptors are CCR1, 3 and 5. The chemokine C-C motif ligand 5 released by human mesenchymal stem cells originating from bone marrow in a paracrine manner is reported to enhance the metastatic capabilities of cancer stem cells (Cuiffo & Karnoub, 2012). One of 7 increased chemokines in solid tumor compared to normal based on TCGA data analysis (Fig. 2).

-In the triple-negative (TNBC) subtype of breast cancer, higher levels of CCL5 than in non-TNBC are pointed out (Hozhabri, Moghaddam, Moghaddam, & Mohammadian, 2022).

-In the study of Pervais, no variation was observed the levels of CCL5 between healthy control, stage II and III patient samples (Pervaiz et al., 2019).

As shown in Figure 2, 3, 4 and Supp. Fig 1, the levels of CCL5 expression tumor and NAT displayed significantly augmented expression compared to samples from healthy breast tissue. This demonstrates its capacity as a potential marker in breast cancer.

According to TCGA data (Fig 5 M-O and Supp. Fig 5E), CCL5 expression levels appear to be higher in tumor samples when compared to normal breast samples. This difference continues to be maintained in cancer stages from stage 1 to stage 3, but it disappears in stage 4, which has normal tissue CCL5 levels. Examining cancer subtypes, the expression levels of this chemokine show an increasing trend from luminal cancer to TNBC cancer subtype, giving a signals of holding to be a potential subtype marker. Survival plot shows a safe 8 years good survival probability for high expressed patients (p<0.05) (Fig 8E).

**Figure 8.**
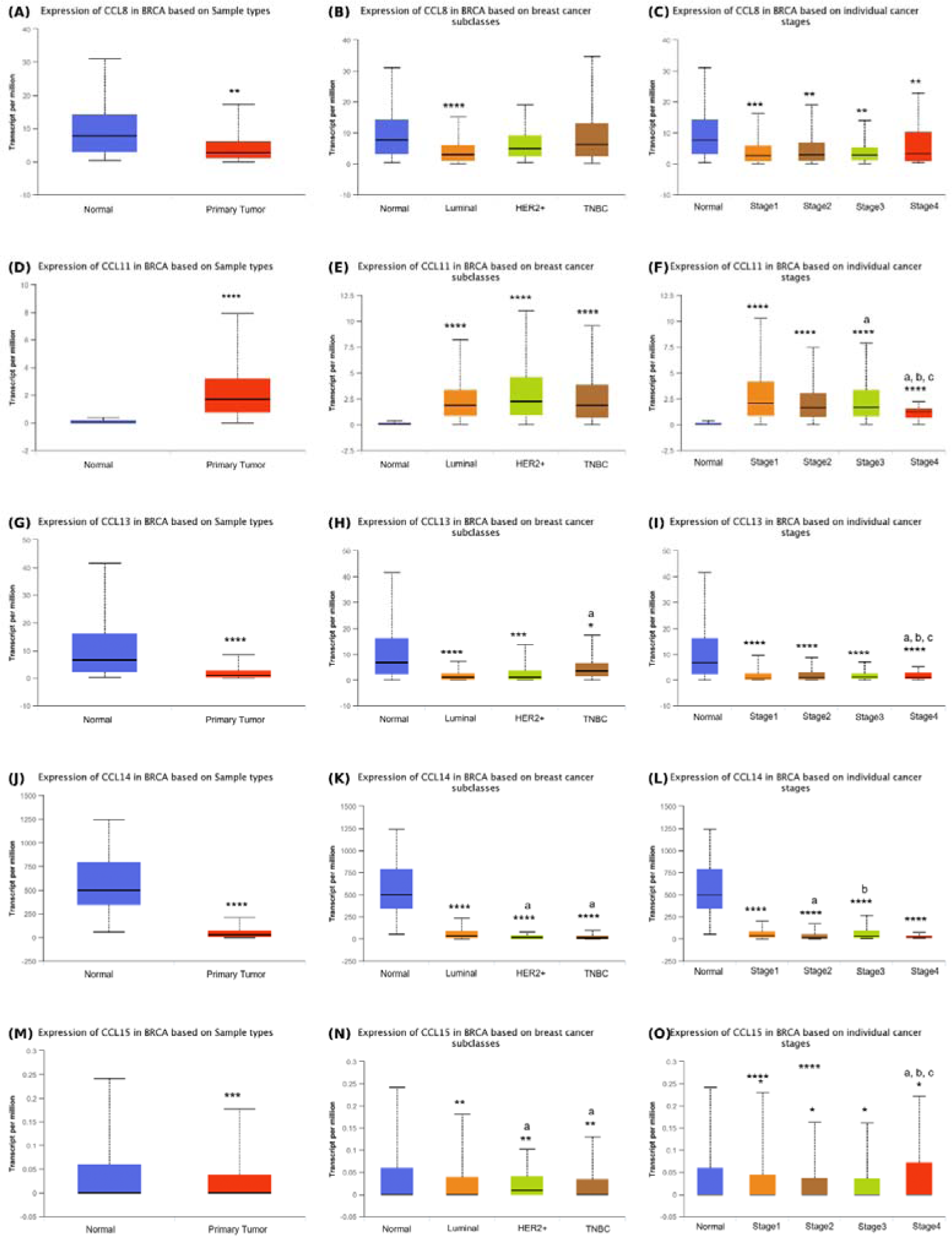
The expression of CCLs 8, 11 and 13-15. *Sample type*: The expressions of CCL8 (**A**), CCL11 (**D**), CCL13 (**G**), CCL14 (**J**) and CCL15 (**M**) in normal (n=114) and breast cancer tissue samples (n=1097). *Breast cancer molecular subclasses*: The expressions of CCL8 (**B**), CCL11 (**E**), CCL13 (**H**), CCL14 (**K**) and CCL15 (**N**) in normal tissue (n=114) and breast cancer subtypes of luminal (n=566), HER2^+^ (n=37) and triple negative (TNBC, n=117). *Cancer stages:* The expressions of CCL8 (**C**), CCL11 (**F**), CCL13 (**I**), CCL14 (**L**) and CCL15 (**O**) in normal tissue and different stages including stage 1 (n=183), stage 2 (615), stage 3(n=247) and stage 4 (n=20). **Note:** In all graphs, *, **, *** and **** symbols designate a significant difference of p<0.5, p<0.01, p<0.001 and p<0.0001, respectively. In the individual cancer stages graph, the small letters of a, b and c designate a significant difference between stages 1, 2 and 3, respectively. In the breast cancer subclasses graph, the small letters of a and b designate a significant difference between luminal and HER-positive subclasses, respectively.

## C-C motif chemokine ligand 7 (CCL7)

Another name for CC7 is MCP-3 and its receptors are CCR1, 2 and 3. Fibroblast-secreted CCL7 is reported to provoke tumor cell growth. CCL2, CCL7 and CCL8 chemokines convey tumor-nurturing properties of fibroblasts with separate impacts on the tumor milieu (Rajaram, Li, Egeblad, & Powers, 2013). One of 7 increased chemokines in solid tumor compared to normal based on TCGA data analysis (Fig. 2).

As shown in Figure 2, 3, 4 and Supp. Fig 1, the levels of CCL7 expression are significantly higher in tumor with no difference between NAT and healthy breast tissues samples, indicating potential of CCL& to be tumor specific marker.

In TCGA database, expression of this chemokine in tumor samples is significantly higher than normal breast tissue. In subclass analysis, there is an obvious ascending expression levels from luminal cancer type to TNBC. Moreover, while the expression is high in all stages of cancer, it decreased to control levels in stage 4. In Kaplan Meier survival curve, although no significance detected, there is a high survival trend in the plot of the patients with high expression of CCL7 when compared to low/moderately expressed ones.

## C-C motif chemokine ligand 8 (CCL8)

Its other name is MCP-2 and CCL8 is a ligand of several receptors: CCR1, CCR2B, CCR5 and CCR8. CCL8 accumulation between epithelium and stroma contributes the breast cancer propagation. Neoplastic epithelial signals elevate CCL8 levels in stromal fibroblasts adjacent to the tumor and the potential metastasized tissues like the lungs and brain. Gradients of chemoattractant, CCL8 determine the fate of the tumor (Farmaki, Chatzistamou, Kaza, & Kiaris, 2016). Farmaki et al. (2016) further speculate that this role of CCL8 is associated with more breast and cervical cancers rather than other cancer types. CCL8 levels are documented to progressively rise during tumor growth and plunge to control levels after tumor remission (Pitteri et al., 2011).

As shown in Figure 2, 3, 4 and Supp. Fig 1, CCL8 expression levels are higher in tumor and NAT tissues relative to healthy breast tissue samples. When compared to tumor and adjacent normal tissues, NAT has a significantly higher value of CCL8 expression. The pronounced presence of CCL8 in NAT points towards its enhanced role within the context of the tumor microenvironment.

According to TCGA data (Fig 6 A-C and Supp. Fig 5G), CCL8 expression in primary tumor expression versus normal tissue is significantly lower. In cancer luminal breast cancer subtype, this low expression is still obvious (p<0.0001). However, this is not apparent in the cancer subclasses of HER2+ and TNBC tumor samples with no significant difference from normal tissue samples regarding CCL8 expression. All stages of cancer also demonstrate a low expression profile of CCL8. Collectively, this dataset showcases that CCL8 lower expression can be pertinent to luminal subtype of breast cancer. There was no difference detected in the survival plots of the patients with high and low/medium expression of CCL8.

## C-C motif chemokine ligand 11 (CCL11)

Its other name is eotaxin. Its receptors include CCR2, CCR3 and CCR5, with the highest affinity to the middle. It selectively attracts eosinophils. In the Wang et al paper, CCL11 is reported to exert a regulatory role in T cells. It augments the frequency of Treg cells and activates the STAT5 signaling pathway, in doing so, fosters immunosuppression and immune tolerance in breast cancer. Therefore, reshaping the immune landscape by blocking CCL11 is a plausible avenue for breast cancer therapy (R. Wang & Huang, 2020). One of 7 increased CC-type chemokines in solid tumor compared to normal based on TCGA data analysis (Fig. 2).

As shown in Figure 2, 3, 4 and Supp. Fig 1, the levels of CCL11 are higher in solid tumor adjacent tissue relative to healthy breast samples. In addition, in solid tumor the expression of CCL11 is almost 2-3 folds more than NAT. CCL11 also shows a strongly indiacative potential of being a solid tumor specific marker.

According to TCGA data (Fig 6 D-F and Supp. Fig 5H), this chemokine shows a disparately clear-cut higher expression values in tumor tissue when compared to normal tissue. This is the only CCL chemokines in TCGA database showing clear cut difference and being expressed at high level in tumor. Most of the obvious differences between tumor and normal tissue regarding CCL chemokine expression are found at low expression of tumor tissue. In this case, this chemokine deserves a special attention as this chemokine expressed at high levels in tumor with chemokines, generally speaking, being notorious to adversely affect the prognosis of cancer. CCL11 levels seem to be significantly higher in all subclasses and stages (all p<0.0001). In addition, despite this significance, there is a trend of stepwise decreasing the expression levels of CCL11 from stage 1 to stage 4 with the latter showing a significant difference from all other 3 stages. No significant survival curve is recorded.

## C-C motif chemokine ligand 13 (CCL13)

Its other name is MCP-4, and its receptors are CCR2 and CCR3 (Fig 5B). A remarkable difference in the expression levels of CCL13 was uncovered between cancer and benign lesions. Breast cancer with a more aggressive course was reported to relate to relatively high levels of CCL13 (Franzen et al., 2019). A strong correlation was detected between Ki67 status and CCL13 levels.

As shown in Figure 2, 3, 5 and Supp. Fig 1, intriguingly, CCL13 levels are lower in solid tumor, but higher in healthy tissue and NAT, with the highest in the latter.

According to TCGA data (Fig 6 G-I and Supp. Fig 5I), this chemokine is also included among the one of the lowest tumor expressing cytokines relative to normal breast tissue with almost 7 -fold of median expression value difference (p=0.0001). It followed a similar trend among each of the 4 stages of cancer with a significant disparity from normal tissue. In molecular subclasses of breast, despite all subclasses exhibiting a significant reduction compared to normal tissue, TNBC subtype also showed a significant difference from luminal subtype with a little bit more increasing trend in expression of CCL13. No significant survival probability was evident with the same high survival trend that is seen in a couple of other chemokines’ survival graphs between the years 5.4 to 10 follow ups.

## C-C motif chemokine ligand 14 (CCL14)

Its other name is HCC-1, and its receptor is CCR1 (Fig 5B).

As shown in Figure 2, 3, 5 and Supp. Fig 1, CCL14 levels are somewhat lower in NAT, but the lowest in solid tumor tissue.

According to TCGA data (Fig 6 J-L and Supp. Fig 5J), this is another chemokine requiring special attention with clear cut difference between tumor or normal tissue expressions. Considering median value expressions, there is a 19-fold difference in-between. This obvious difference is maintained in all stages and molecular types of breast tumor. Therefore, it is conceivable to say that this difference is a subtype and stage free, suggesting CCL14 to have a potential to be marker of breast cancer. First 10 years of survival plot, a better survival data dominates in the patients with high expression of CCL14. Nevertheless, the p values are greater than 0.05 (p=0.059).

## C-C motif chemokine ligand 15 (CCL15)

Its other names are HCC-2, MIP-5, MIP-1-d and Lkn-1. Its receptors are CCR1 and 3 (Fig 5B). According to our TCGA analysis, it is one of the 5 CC-motif chemokines with no significant change between solid tumor and normal breast tissue (Fig. 2).

As shown in Figure 2, 3, 5 and Supp. Fig 1, CCL15 levels are lower and consistent in solid tumor and NAT when compared to healthy breast tissue.

According to TCGA data (Fig 7 M-O and Supp. Fig 5K), TPM values are very low when compared to other chemokines. Although visually on the graphs, significant difference is not obvious, there is a significant diminish in the CCL15 levels of tumor tissue. Other than that, in cancer stage-based analysis, a prominent increase in stage 4 is recorded in dataset. In survival curve, despite a high survival probability between starting after 5 years of surveillance at high expression group, no significant difference was detected between in patients high and low/medium expression of CCL15.

## C-C motif chemokine ligand 16 (CCL16)

Another name for this chemokine is HCC-4. Its receptor is CCR1.

As shown in Figure 2, 3, 5 and Supp. Fig 1, CCL16 levels are lower in NAT when compared to normal breast tissue samples, but the lowest in solid tumor tissue.

According to TCGA data (Fig 5 A-C and Fig 8L), there is an obvious difference in CC16 expression regarding tumor samples versus normal breast samples and this significant difference is also clearly observed in each subclass and each stage. Overall survival graphs of high and low/medium expressions of CCL16 follow a similar trend. In contrast, Shen et al (2021) reported that CCL16 is highly expressed in breast tumors and associated with the progression of the tumor and silencing CCL16 in mice xenografts suppresses tumorigenicity and shrinks the tumor size, decreasing stemness and chemoresistance (Shen et al., 2021). Also, CCL16 is postulated to have anti-cancer effects and to be used in therapy with its recruitment of CD4+ CD8+ and DC cells (Korbecki et al., 2020).

## C-C motif chemokine ligand 17 (CCL17)

Chemokine ligand 17 (CCL17) is also known as thymus-and activation-regulated chemokine (TARC), and its only known receptors are CCR4. CCL17 is mentioned in the paper of Mollica Poeta et al. (2019) that it directly gathers regulatory T cells and TH2 lymphocytes and exacerbate tumor outcome (Mollica Poeta et al., 2019). In the original paper of …, this effect is indicated to not be as effective as CCL22 shows (Gobert et al., 2009). One of 7 increased chemokines in solid tumor compared to normal based on TCGA data analysis (Fig. 2).

As shown in Figure 2, 3, 5 and Supp. Fig 2, CCL17 levels are significantly higher in NAT when compared to normal breast tissue samples and highest in solid tumor tissue (vs both NAT and normal tissue).

According to TCGA data (Fig 7 D-F and Supp. Fig 6A), expression of CC17 in tumor samples are significantly higher than normal breat tissue sampkes. When considering molecular subtypes, namely luminal, her2+ and tnbc. All subtypes possesses a significant high values than normal breast tissue but there is no significant difference among them, despite HER2+ and TNBC subtypes having higher median values. In terms of stage analysis, all three stages (1, 2, and 3) show significantly higher CC17 expression compared to normal samples. However, in the last stage (stage 4), there is no significant difference in CC17 expression. Overall, CCL17 could be a potential marker excluding stage 4, having consistent high expression in tumor samples. CC17 is not specific to subtype or advanced stages on account of the lack of significant differences between subtypes and the absence of a difference in expression in stage 4 tumors. In survival curves, no significant value has been detected. However, a good survival output is visible between the follow-up of 5.5 years to 10, after which an abrupt fall starts in the graph (p>0.05).

## C-C motif chemokine ligand 18 (CCL18)

CCL18 is also called AMAC-a or MIP-4. Its receptor is yet to be known. It is the smallest known human CC-motif chemokine with 89 aa and a molecular weight of 9,849 kDa. Abundant CCL18 expression by tumor-associated macrophages in blood or cancer stroma has been linked to metastasis and shortened patient survival. Through facilitating integrin clustering and inducing their adherence to the extracellular matrix, CCL18 secreted by breast TAMs make cancer cells more invasive. Based on Chen’s 2011 research, PITPNM3 orchestrates the CCL18 effect and activates intracellular calcium signaling. These malignant effects of CCL18 including the invasion and metastasis of breast cancer xenografts can be abrogated by PITPNM3 suppression (J. Chen et al., 2011).

CLL18 (AMAC-1) exists naturally in alveolar macrophages, to a lesser extent in monocyte-derived dendritic cells. CCL18 is released by alternatively activated macrophages (M2 macrophages) through Th2-associated cytokines in vitro. Both these macrophages and alveolar macrophages are implicated in Th1-associated immune reaction suppression (Kodelja et al., 1998). The effect of CCL18 general in cancer is controversial Given that M2 macrophages support tumor invasion and metastasis in breast cancer, and M1 macrophages do the opposite by killing the cancer cells, CCL18 needs closer attention.

As shown in Figure 2, 3, 5 and Supp. Fig 2, CCL18 levels are higher in solid tumor tissue when compared to normal breast tissue samples, but the highest in NAT when compared to both tumor and normal breast tissue.

As shown in Figure 2, 3, 4 and Supp. Fig 1, CCL18 expression increases significantly both in tumor and NAT when compared to normal breast tissue. When compared to tumor and adjacent normal tissues, it’s noticeable that NAT exhibits elevated levels of CCL18 expression. The elevated expression of CCL18 in NAT may imply a pivotal role within the context of the tumor microenvironment. It has the potential to be utilized as a marker for diagnosing breast cancer, given its significant high impression in tumor tissue and its environment. Let’s also check its trend in terms of cancer subtype and stages below.

CCL18 expression is lower in tumor tissues with less variation when compared to normal tissue in breast (Fig 7 G-I and Supp. Fig 6B). Low expression of CCL18 might be sub-type specific. Luminal subtype expression is significantly lower than both normal tissue and TNBC subtype, despite no difference with HER2+ subtype but still having lower median value. Also, it has decreased expressions in all stages with lowest median value and variation in the last stage (Fig 7I). In the overall Kaplan Meier survival plot, a high survival outcome starts almost after 5 years at high expressions of CCL18; however, no significant p value is detected between graphs (Supp. Fig 6B).

## C-C motif chemokine ligand 19 (CCL19)

-CCL19 (MIP3β or exodus-3) binds to receptor CCR7 and is mostly expressed in T cell zones like the thymus and lymph nodes.

As shown in Figure 2, 3, 5 and Supp. Fig 2, there is no difference between the expression levels of healthy breast, solid tumor and NAT.

According to the bioinformatics study of Wang et al. who harnessed the TCGA database, CCL19 overexpression is reported to show a better prognosis by communicating infiltrating immune cells and modulating tumor microenvironment (J. Wang et al., 2022), In conjunction with CCL21, CCL19 is recorded to possess a principal role in the interplay between dendritic cells and naïve T cells and activate TH1-polarized immunogenicity (Nguyen-Hoai et al., 2016). According to our TCGA analysis, it is one of the 5 CC-motif chemokines with no significant change between solid tumor and normal breast tissue (Fig. 2).

First, there is a meaningful difference between normal and tumor tissue expression levels of CCL19 (p<0.0001). According to our interpretation from TCGA overall survival graph of CCL19 (p>0.05), although first 11 years of follow-up demonstrate an obvious affirmative effect of CCL19 on patients, p value is insignificant. Furthermore, there is no clear change trend in stage-based expression analysis with significant high values in the first 3 stages relative to normal tissue (Fig 7 J-L and Supp. Fig 6C).

## C-C motif chemokine ligand 20 (CCL20)

Other names of CCL20 are exodus-1 and MIP-3α and its receptor is CCR6. CCL20 expression is higher in breast tumor samples than normal tissue according to TCGA data (Fig 7M).

As shown in Figure 2, 3, 5 and Supp. Fig 2, the levels of CCL20 expression are significantly higher in tumor with no difference between NAT and healthy breast tissues samples, indicating potential of CCL20 to be tumor specific marker.

One of 7 increased chemokines in solid tumor compared to normal based on TCGA data analysis (Fig. 2). CCL20 expression does not exhibit a significant overall survival analysis. In molecular subtypes, there is an ascending expression trend from luminal type to TNCB, severe type of breast cancer but not a clear stage-specific trend. ((Fig 9 M-O and Supp. Fig 6D).

**Figure 9.**
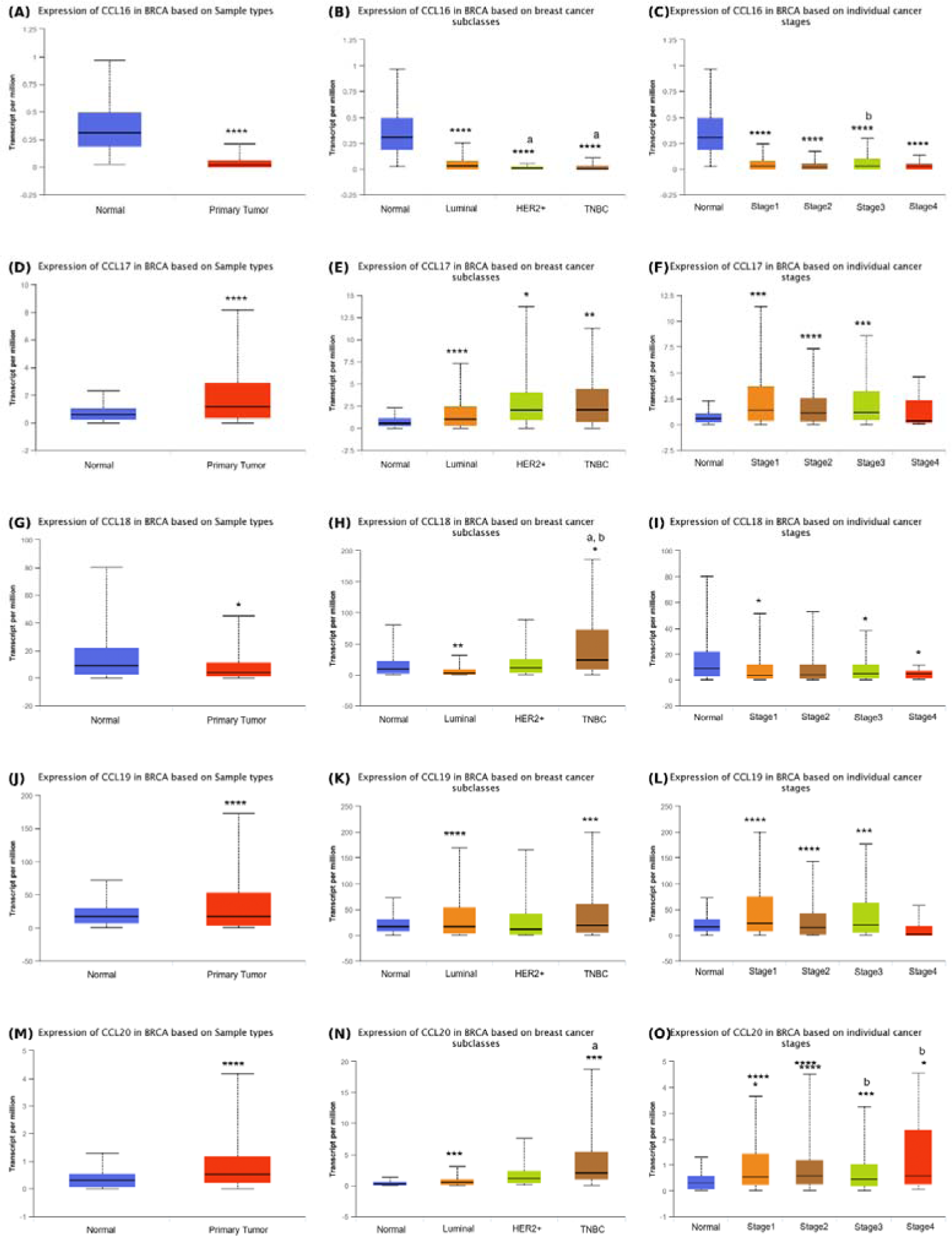
The expressions of CCLs 16-20. *Sample type*: The expressions of CCL16 (**A**), CCL17 (**D**), CCL18 (**G**), CCL19 (**J**) and CCL20 (**M**) in normal (n=114) and breast cancer tissue samples (n=1097). *Breast cancer molecular subclasses*: The expressions of CCL16 (**B**), CCL17 (**E**), CCL18 (**H**), CCL19 (**K**) and CCL20 (**N**) in normal tissue (n=114) and breast cancer subtypes of luminal (n=566), HER2^+^ (n=37) and triple negative (TNBC, n=117). *Cancer stages:* The expressions of CCL16 (**C**), CCL17 (**F**), CCL18 (**I**), CCL19 (**L**) and CCL20 (**O**) in normal tissue and different stages including stage 1 (n=183), stage 2 (615), stage 3(n=247) and stage 4 (n=20). **Note:** In all graphs, *, **, *** and **** symbols designate a significant difference of p<0.5, p<0.01, p<0.001 and p<0.0001, respectively. In the individual cancer stages graph, the small letters of a, b and c designate a significant difference between stages 1, 2 and 3, respectively. In the breast cancer subclasses graph, the small letters of a and b designate a significant difference between luminal and HER-positive subclasses, respectively.

## C-C motif chemokine ligand 21 (CCL21)

It is also called 6Ckine or Exodus-2 and it binds to CCR7 as a receptor. CCL21 is suggested as a potential prognostic marker associated with immune cell infiltration, carcinogenesis and tumor immune escape (B. Chen et al., 2020). A shorter OS and RFS are recorded in low-expressed CCL21 BC patients, which is proposed to increase immunogenicity (Hozhabri et al., 2022). In conjunction with CCL19, CCL21 is documented to take a vital part in establishing a bridge between dendritic cells and naïve T cells and activates TH1-polarized immunogenicity (Nguyen-Hoai et al., 2016).

As shown in Figure 2, 3, 6 and Supp. Fig 2, while there is no difference between normal breast and tumor adjacent breast samples, tumor samples follow a significant reduced levels of CCL21 expression.

According to TCGA data, a clear-cut disparity is observed between the CCL21 levels of tumorigenic and normal breast tissue samples, underlining the remarkable potential and inherent value of this chemokine in breast cancer. This striking difference (almost 13-fold difference between median values) is also visible across every sub-groups when categorizing the cancer samples into subclasses and stages. It seems to be independent from context-based situations. (Fig. 10A-C and Supp. Fig 6E). Last but not least, high expression of CCL21 in breast cancer patients also offers an unambiguous overall survival plot (Supp. Fig 6E, p=0.015).

**Figure 10.**
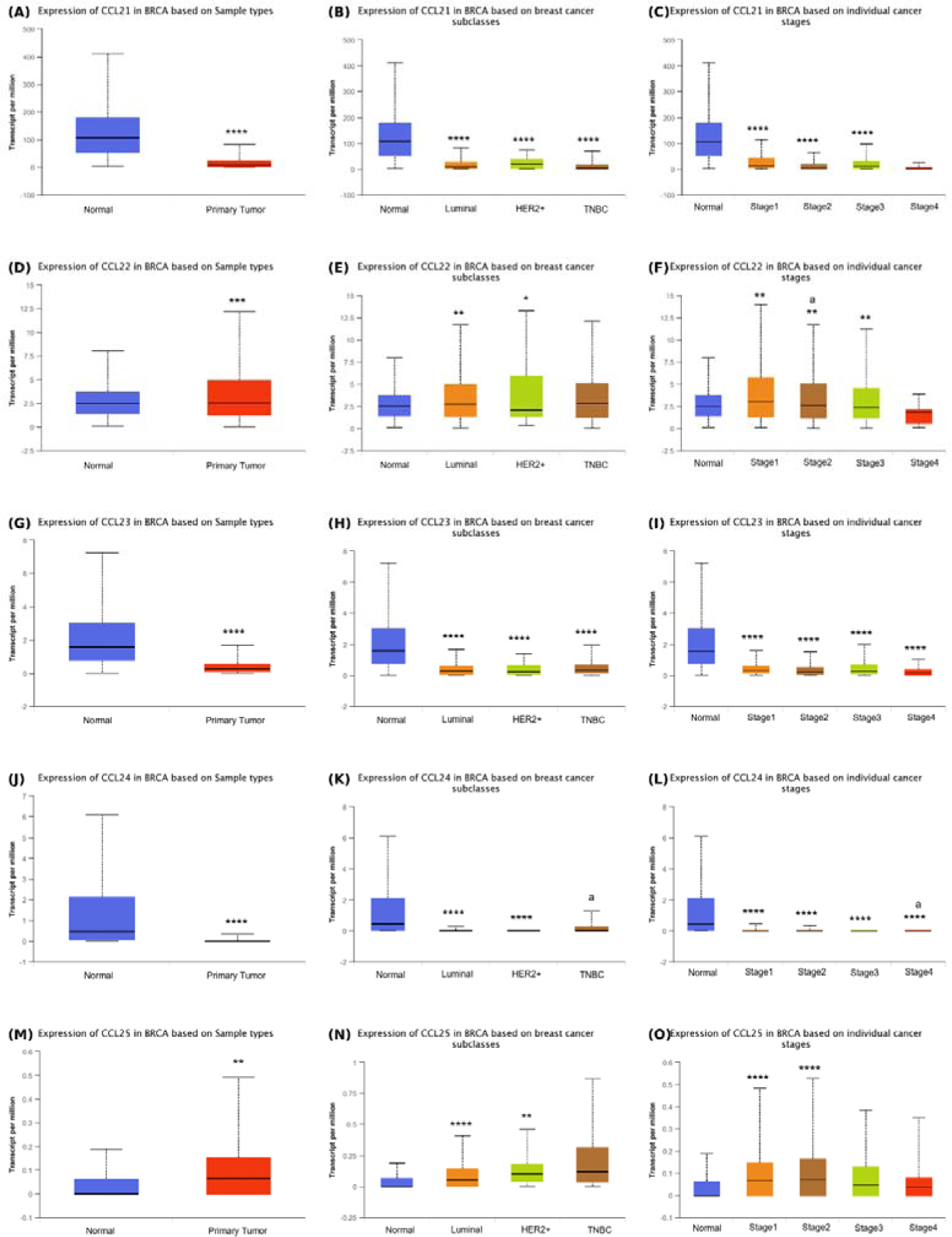
The expression of CCLs 21-25. *Sample type*: The expressions of CCL21 (**A**), CCL22 (**D**), CCL23 (**G**), CCL24 (**J**) and CCL25 (**M**) in normal (n=114) and breast cancer tissue samples (n=1097). *Breast cancer molecular subclasses*: The expressions of CCL21 (**B**), CCL22 (**E**), CCL23 (**H**), CCL24 (**K**) and CCL25 (**N**) in normal tissue (n=114) and breast cancer subtypes of luminal (n=566), HER2^+^ (n=37) and triple negative (TNBC, n=117). *Cancer stages:* The expressions of CCL21 (**C**), CCL22 (**F**), CCL23 (**I**), CCL24 (**L**) and CCL25 (**O**) in normal tissue and different stages including stage 1 (n=183), stage 2 (615), stage 3(n=247) and stage 4 (n=20). **Note:** In all graphs, *, **, *** and **** symbols designate a significant difference of p<0.5, p<0.01, p<0.001 and p<0.0001, respectively. In the individual cancer stages graph, the small letters of a, b and c designate a significant difference between stages 1, 2 and 3, respectively. In the breast cancer subclasses graph, the small letters of a and b designate a significant difference between luminal and HER-positive subclasses, respectively.

## C-C motif chemokine ligand 22 (CCL22)

It is known with multiple different names including MDC and STCP-1 and it binds to CCR4 as a receptor. In the study where 199 invasive breast cancer tissues were stained immunohistochemically, CCL22 is expressed in the majority of breast cancer tissues. Lower-grade tumors indicated high levels of CCL22 (Kuehnemuth et al., 2018). Consistently, there is higher expression of CCL22 in TCGA data at early stages of tumor, considering that tumor grade is parallel to stage progression (See Fig 8F. In addition, in the study, no correlation between regulatory T cells and CCL22 was detected, whereas; on the contrary, CCL22 is suggested to attract tumor-infiltrating regulatory T cells through CCR4 in the study of Gobert et al. (2009). According to our TCGA analysis, it is one of the 5 CC-motif chemokines with no significant change between solid tumor and normal breast tissue (Fig. 2).

As shown in Figure 2, 3, 6 and Supp. Fig 2, there is an increasing trend of expression of CCL22 with the order of normal breast samples-NAT-tumor samples.

In the TCGA data (Fig 10 D-F, and Supp. Fig 6F), stage-based decrease of CCL22 is highly evident in breast tumor samples. There is a descending trendline in the median values of CCL22 expression from stage 1 to stage 4 (median values: 3.0, 2.6, 2.4, 1.9 transcripts/million, respectively) with significant difference of first 3 stages compared to normal tissue expressions. In stage 4, despite not statistically meaningful, CCL2 levels are even lower than those in normal tissue. However, the other 3 states have significantly high levels of CCL2 with orderly decreasing trend. It is also worth a subclasses-based difference between expressions of normal tissue versus luminal and HER2+ subtypes but not TNBC.

## C-C motif chemokine ligand 23 (CCL23)

It is also called MIP-3 or MPIF-1 and it binds to CCR1 as a receptor. A linkage between the aggressiveness of breast cancer with relatively high levels of CCL23 is documented in the study of Franzen et al. (2019).

As shown in Figure 2, 3, 6 and Supp. Fig 2, CCL23 levels are lower in NAT when compared to normal breast tissue samples, but the lowest in solid tumor tissue.

This particular chemokine has a significantly lower values relative to normal tissue levels in all molecular types of tumors and cancer stages. It does not have any significant Kaplan Meier plot results (Fig. 10G-I and Supp. Fig 6G). It is not redundant to say that 6-fold difference between median expression values and having less overlapping values between the normal and cancer samples could propose CCL23 to serve a future potential breast cancer marker (Fig 10C).

## C-C motif chemokine ligand 24 (CCL24)

It is also called MPIF-2 or eotaxin-2 and its receptor is CCR3. An elevated level of CCL24 was related to worse OS in BC patients (Hozhabri et al., 2022).

As shown in Figure 2, 3, 6 and Supp. Fig 2, intriguingly like CCL13 levels, CCL24 levels are lower in human solid tumor, but higher in healthy tissue and NAT, with the highest in the latter. Lower levels of CCL24 in tumor (p<0.0001, Fig. 10J) is not a stage dependent with all stages of cancer maintaining low levels of expression. In overall survival analysis, no significant correlation was detected with the alteration of expression rate. (Fig 10 J-L and Supp. Fig. 6H).

## C-C motif chemokine ligand 25 (CCL25)

It is also called TECK (Thymus expressed chemokine) and and its receptor is CCR9. It is the biggest human CC-motif chemokine with 150 aa and 16,639 kDa. Its receptor is CCR9. One of 7 increased chemokines in solid tumor compared to normal based on TCGA data analysis (Fig. 2).

As shown in Figure 2, 3, 6 and Supp. Fig 2, the levels of CCL25 expression are significantly higher in tumor with no difference between NAT and healthy breast tissues samples, indicating potential of CCL25 to be solid tumor specific marker.

According to TCGA dataset (Fig. 10), tumor samples show a significantly higher expression of CCL25 chemokine. Furthermore, CCL25 expression increases prominently in the first 2 stages of cancer but decreases to normal tissue levels at stage 3 and 4 in orderly manner. Based on molecular subclasses of breast cancer, in luminal and HER2+ subtypes present a significant high level of this chemokine (p<0001 and p<0.01, respectively) with no significant difference in TNBC. When we look at the survival analysis of low and high expressed patients (p=0.02, Supp. Fig. 6K), CCL25 exhibits a biphasic survival graph. While a better survival is visible up to 9-10 years in the early period in patients with high expression of CCL25, a sharp drop occurs in survival rates after 20 years onwards. This sharp step decrease may not be accurate. This may be because of the decreasing number of patients at the far right of a survival curve, which needs to be estimated cautiously. This applies to all survival curves discussed.

Drawn from these results, we can plausibly infer that high CCL25 shows beneficial effects at the early stages of breast cancer; however, at late stages it commences to exert adverse effects. Collectively, this duality underscores the relevance of the context-dependent or stage-specific intervention in breast cancer.

## C-C motif chemokine ligand 26 (CCL26)

It is also known as Eotaxin3 or macrophage inflammatory protein 4-alpha (MIP-4-alpha) and its receptor is CCR3.

As shown in Figure 2, 3, 6 and Supp. Fig 2, CCL26 levels are lower in both solid tumor and NAT tissues when compared to normal breast samples. In addition, solid tumor samples mean is a little higher than the NAT.

It is one of the two CC-motif chemokines that does not exhibit a statistically significant difference between the normal and tumorigenic breast tissue according to UALCAN database and our TCGA analysis (Fig. 11A-C). Likewise, the comparison of overall survival between different CCL26 expression level patients did not result in any significant difference (Supp. Fig 6C).

**Figure 11.**
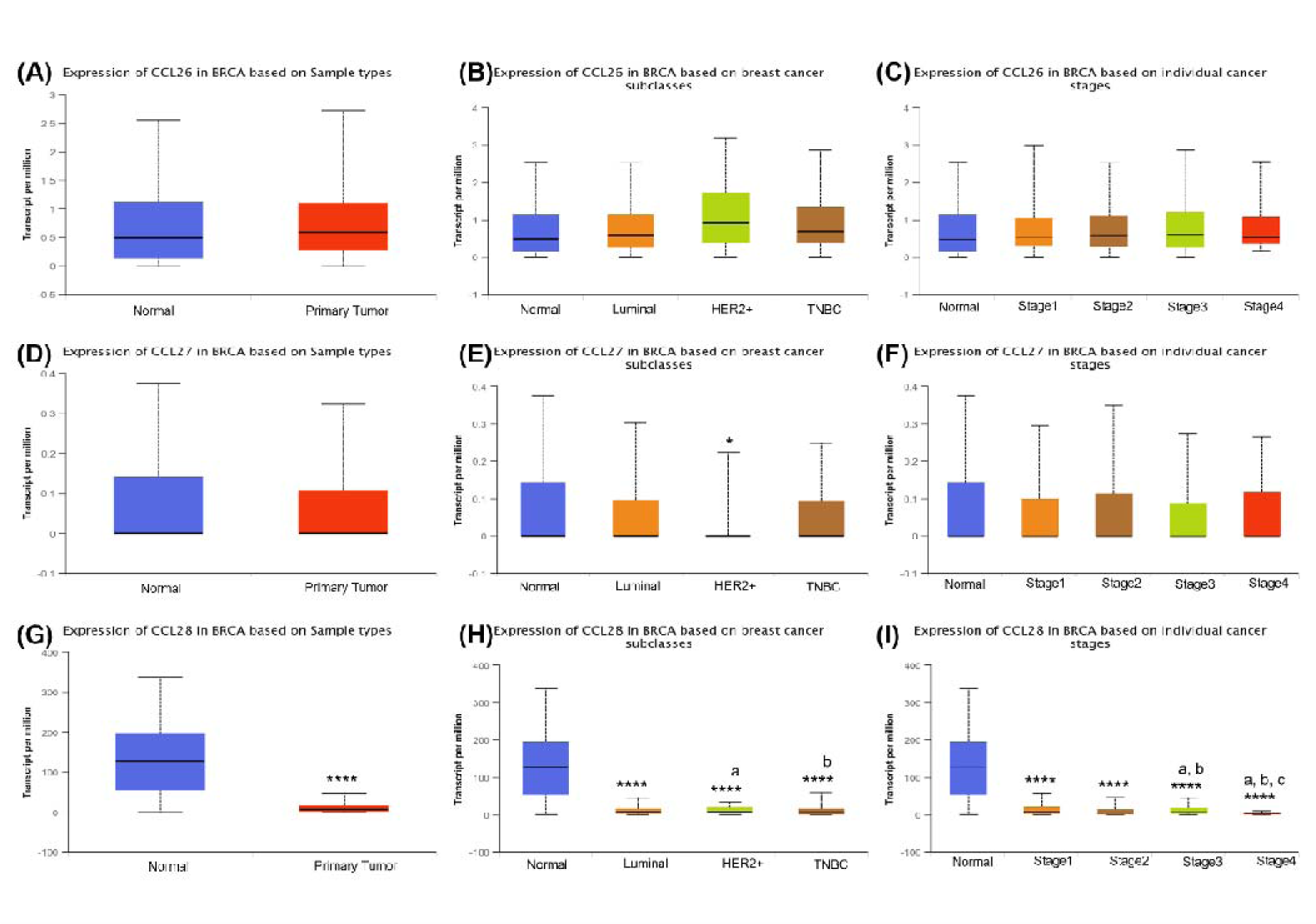
The expression of CCLs 26-28. *Sample type*: The expressions of CCL26 (**A**), CCL27 (**D**) and CCL28 (**G**) in normal (n=114) and breast cancer tissue samples (n=1097). *Breast cancer molecular subclasses*: The expressions of CCL26 (**B**), CCL27 (**E**) and CCL28 (**H** in normal tissue (n=114) and breast cancer subtypes of luminal (n=566), HER2^+^ (n=37) and triple negative (TNBC, n=117). *Cancer stages:* The expressions of CCL26 (**C**), CCL27 (**F**) and CCL28 (**I**) in normal tissue and different stages including stage 1 (n=183), stage 2 (615), stage 3(n=247) and stage 4 (n=20). **Note:** In all graphs, *, **, *** and **** symbols designate a significant difference of p<0.5, p<0.01, p<0.001 and p<0.0001, respectively. In the individual cancer stages graph, the small letters of a, b and c designate a significant difference between stages 1, 2 and 3, respectively. In the breast cancer subclasses graph, the small letters of a and b designate a significant difference between luminal and HER-positive subclasses, respectively.

## C-C motif chemokine ligand 27 (CCL27)

CCL28 is also known with multiple different names including Eskine and ILC and its only known receptor is CCR10 (Fig 2B).

As shown in Figure 2, 3, 6 and Supp. Fig 2, CCL27 expression levels have a lower trend of expression when compared to normal breast tissue.

It is one of the two CC-motif chemokines that does not exhibit a statistically significant difference between the normal and tumorigenic breast tissue according to UALCAN database and our TCGA analysis. (Fig. 11A-C). Intriguingly, according to Kaplan Meier analysis on breast cancer patients with higher (n=272) and lower or medium (n=809) CCL27 expressions, the patients with higher expression of CCL27 demonstrated a longer survival up to 16.5-year follow-up (See Supp. Fig. 6L, p<0.05). This brings up the question of whether exogenous or induced CCL27 can exert an antiproliferative effect on breast cancer.

## C-C motif chemokine ligand 28 (CCL28)

CCL28 is also known as mucosae-associated epithelial chemokine (MEC), CCK1 and SCYA28, and its receptors are CCR3 and 10 (Fig 2B). CCL28 is a mucosa-associated epithelial chemokine and ligand for CCR3/CCR10. Regarding its impact on breast cancer, there are contentious results. According to the Northern blot results of Craig et al 2001, no or very low CLL28 expression was found in most human breast tumors relative to the adjacent normal tissue. Likewise, in situ hybridization experiment indicated that CCL28 was abundantly expressed in the healthy mammary ductal epithelium. However, its expression was either absent or notably reduced in different types of epithelial-derived mammary tumors (Mickanin, Bhatia, & Labow, 2001). On the other hand, the results of Yang et al (2017) are in a different trend. In their study, CCL28 overexpressing cells were observed to promote tumor growth and cellular proliferation and boost the migratory, invasive and metastatic characteristics in breast cancer both *in vivo* and *in vitro* by MAPK-mediated anti-apoptosis (Yang, Liu, Lin, Shi, & Ou, 2017).

As shown in Figure 2, 3, 6 and Supp. Fig 2, like CCL13 and CCL24 levels, CCL28 levels also follow a lower expression in solid tumor, but higher in healthy tissue and the highest in NAT.

According to data retrieved from TCGA (See Fig. 11), the levels of chemokine CCL28 are at very low levels in primary tumor samples when compared to normal tissue thereof, regardless of breast cancer type and specific cancer stages with stage 4 the lowest (Fig. 11G-I).

All in all, it is conceivable that CCL28 levels plunge in breast cancer tissue. Given Yang et al’ study that CCL28 expression encourages tumor development, we expect high CCL28 expression in the tumor. However, this is not the case. Therefore, we think this suppression can be a sort of homeostatic mechanism in the affected tissue site or because already other chemokines with similar outcomes are active, CCL28 remains inactive.

## Collective results of CCLs

In the study of B. Chen et al. (2020), mRNA expression levels of CCLs 2/3/4/5/7/8/11/17/19/20/22 in breast cancer tissue were significantly upregulated, on the other hand, the mRNA expression levels of CCL14/21/23/28 were downregulated when compared to normal tissues. In addition, in cancer tissue, methylation was found to diminish the expression of CCLs 2/5/15/17/19/20/22/23/24/25/26/27. Moreover, a correlation was highlighted between low expression of CCL3/4 and 23 and the development of drug resistance in breast cancer. Low CCL19/21 and 22 and high CCL8 were suggested to link with a poor prognosis. According to survival data analyzed employing the Kaplan-Meier plotter, breast cancer patients with higher mRNA levels of CCL8/18 and lower mRNA levels of CCL1/3/4/5/11/13/15/16/19/21/22/25/27 portrayed a significantly poorer recurrence-free survival (RFS) outcome. Finally, they presented cytokines CCL8 and CCL21 as putative prognostic markers possibly associated with resistance to chemotherapy, cancer initiation and immune escape.

In the study of Thomas et al. (2019), in breast cancer tissues, they detected an excessive expression of CCL5, −7, −11, −17, −20, −22 and −25. They proposed that high mRNA levels of CCL7, −8, −17, −20 and −25 are indicative of a reduced OS. CCL7 and CCL8 were suggested to be linked with decreased relapse-free survival (Thomas, Mir, Kapur, Bae, & Singh, 2019).

In the bioinformatics study of Hozhabri et al. (2022), while longer OS and RFS was found to be linked with elevated expression levels of CCL4, 5, 14, 19, 21 and 22 w, high level of CCL24 was associated with worse OS in BC patients. Moreover, the results of CCL5, CCL8, CCL14, CCL20, and CCL27 were prominently parallel with the clinical outcomes of BC patients.

## The expression changes of CC-type chemokine receptors

As illustrated in Supp. Fig 3, 4 and 7, The expressions of seven CCR including CCRs 1-6 and 8 are significantly higher in tumor tissue and NAT when compared to healthy breast tissue. Conversely, the expressions of CCRs 9 and 10 are higher in healthy breast tissue relative to both tumor tissue and NAT. Regarding CCR7, the expression in tumor and normal breast samples are similar but lower in NAT.

## Concluding Remarks

To wrap up, herein, we presented the gene expression signatures of chemokines in tumor milieu and general individual CC-motif chemokine-based information, utilizing TCGA, UALCAN and Xena UCSC websites with literature search. In this study, we presented, organized, edited different websites human breast cancer data, and analyzed ourselves, providing an easy way for researcher to check the trends of CC-type chemokines and their receptors with clear illustrational diagrams regarding their differential expressions. Chemokines, more broadly, are very crucial in the initiation, inhibition as well as development and growth of tumors. They are not only attracting the miscellaneous sort of cells in site but also modulating or switching between the cellular states and they act as a gradient manner. Increasing our knowledge about these promiscuous cytokines interaction and function will open a flexible and alternative avenue for cancer therapy.

## Data availability

Source codes used in this article are accessible via github link below: https://github.com/sedatkacar56/xenadataprepare.md.git. Any additional information required reported in this paper is available from the corresponding author.

## ACKNOWLEDGEMENTS

Gene expression (RNAseq) data were downloaded from UALCAN (http://ualcan.path.uab.edu/analysis.html). The results shown here are in whole or part based upon data generated by the TCGA Research Network: https://www.cancer.gov/tcga.

---

**Supp. Fig 1.**
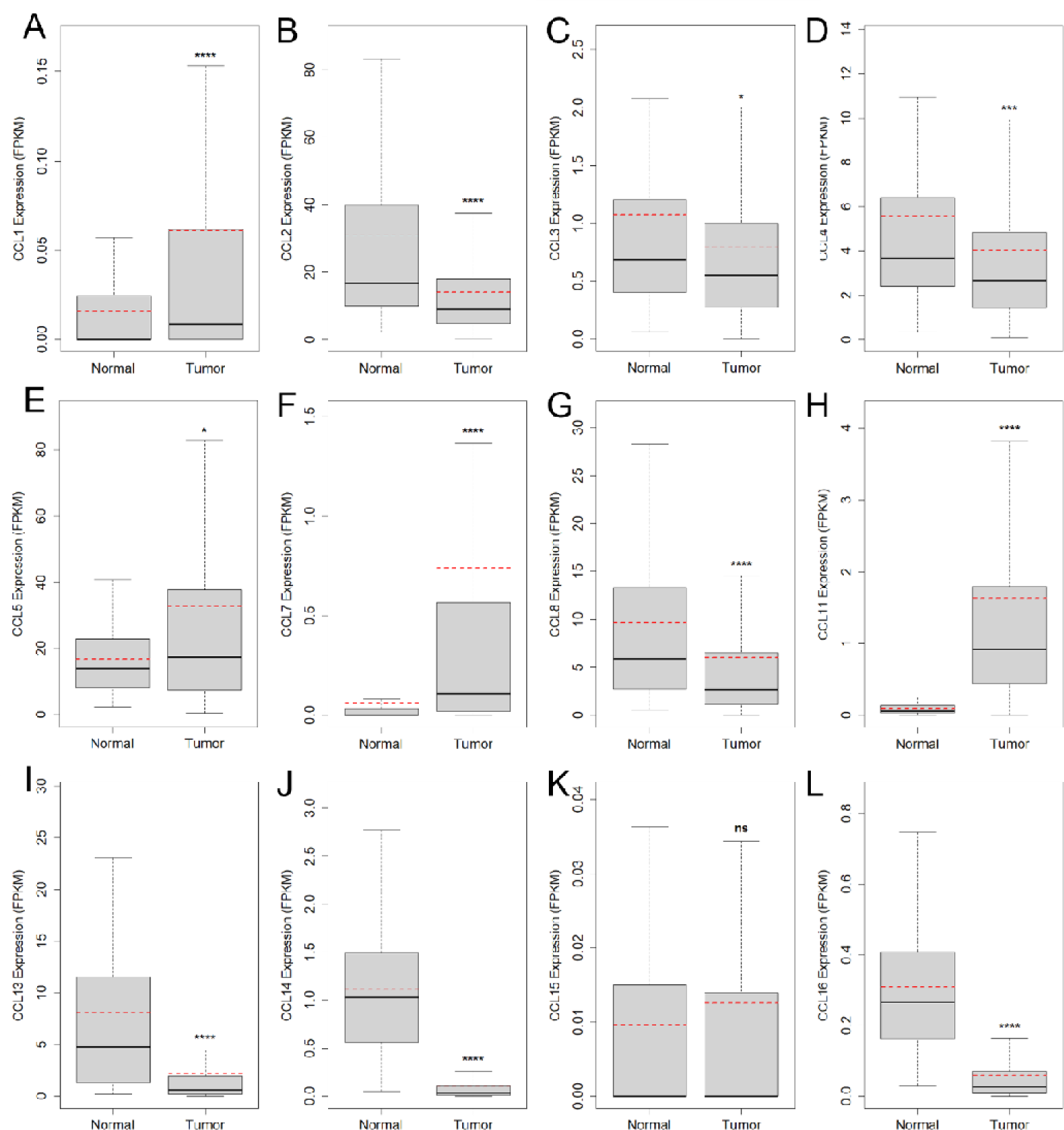
The expressions of CCLs 1-5, 8, 11, 13-16 retrieved from TCGA database. database (n=88 for normal solid tissue, n=104 for tumor)**. Note:** In all graphs, *, **, *** and **** symbols designate a significant difference of p<0.5, p<0.01, p<0.001 and p<0.0001, respectively based off of either t-test or Mann Whitney test.

**Supp. Fig 2.**
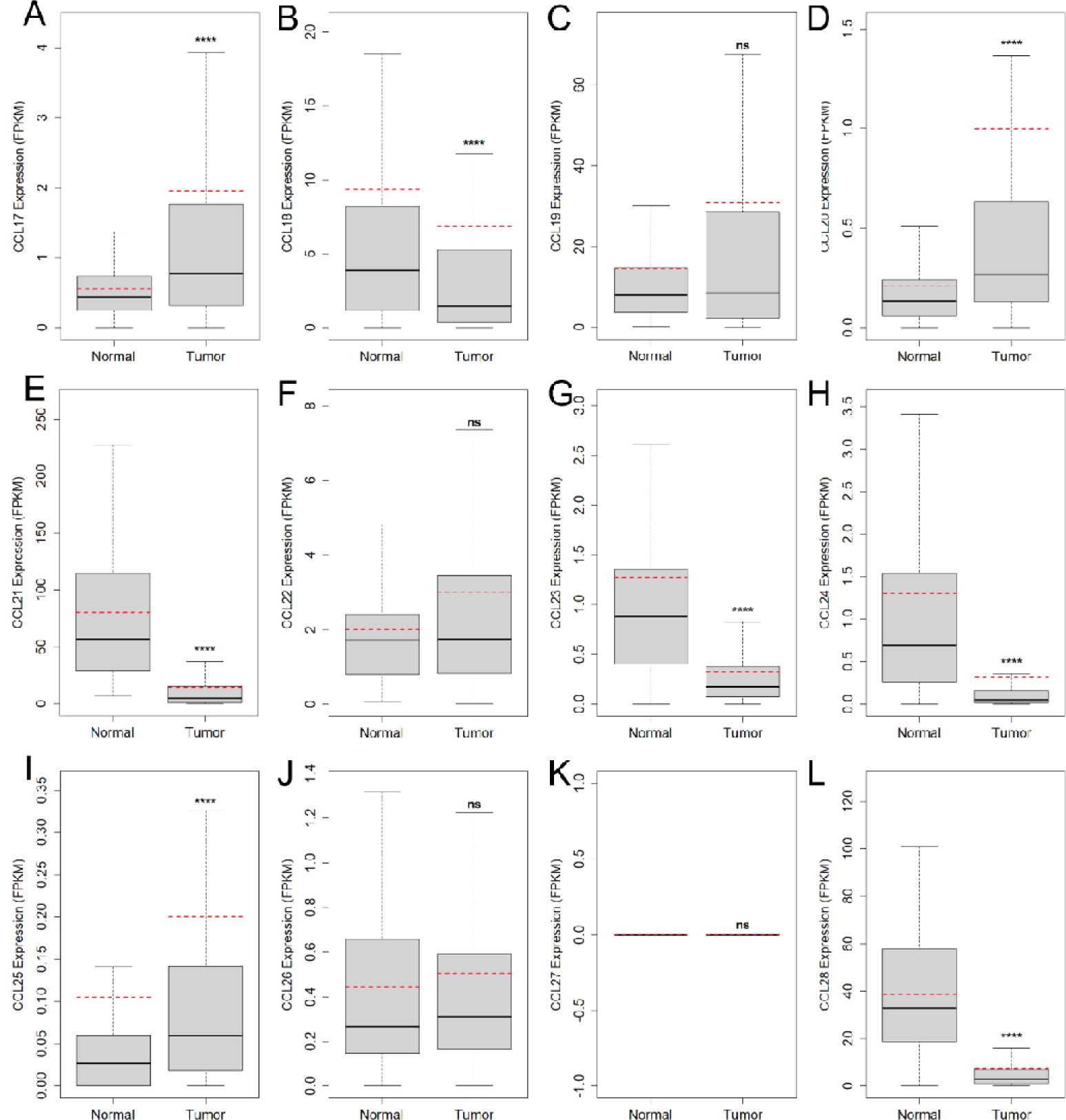
The expressions of CCLs 16-28 retrieved from TCGA database (n=88 for normal solid tissue, n=104 for tumor) **Note:** In all graphs, *, **, *** and **** symbols designate a significant difference of p<0.5, p<0.01, p<0.001 and p<0.0001, respectively based off of either t-test or Mann Whitney test.

**Supp. Fig 3.**
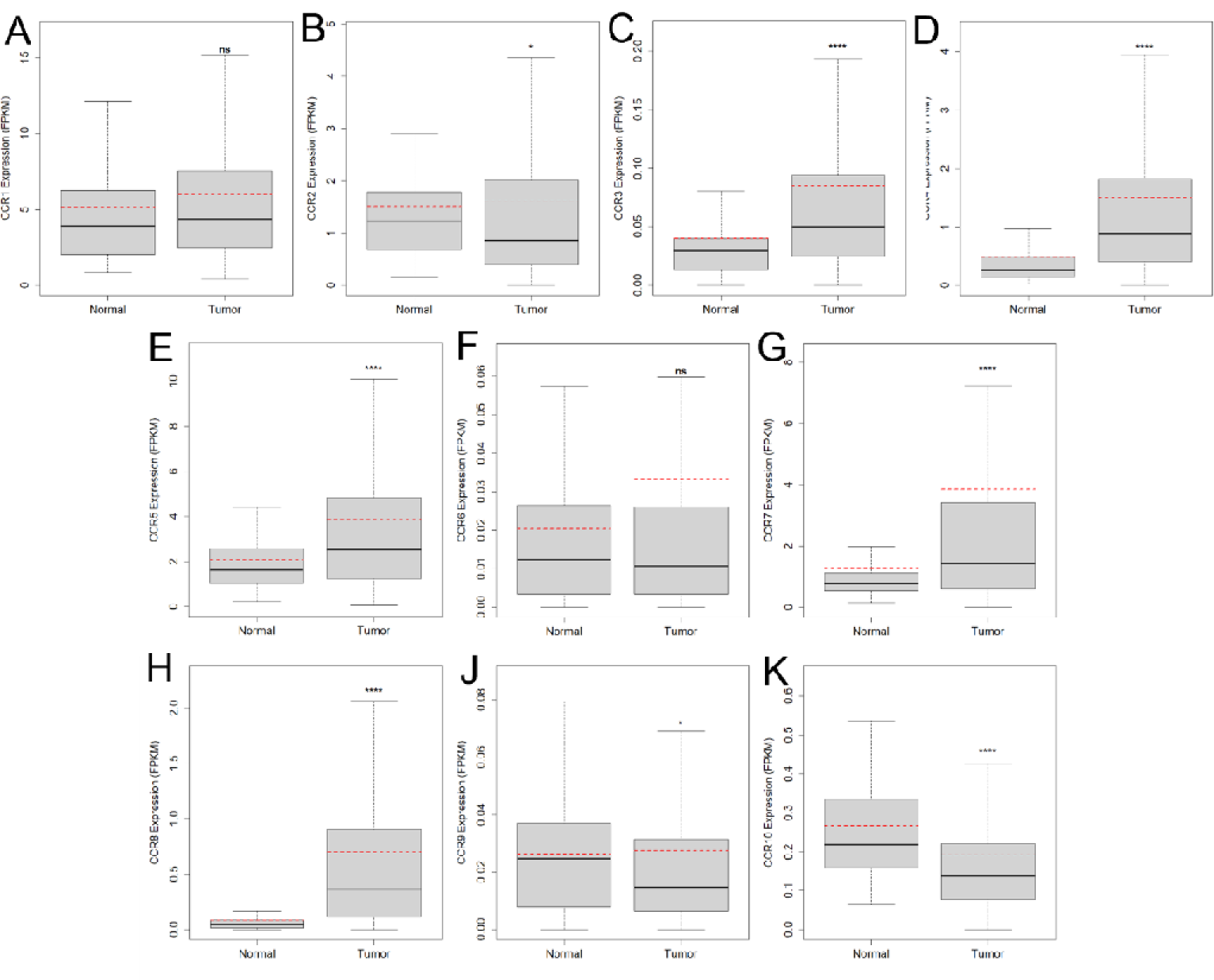
Expression levels of CC Chemokine Receptors, CCRs 1-10 in tumor versus normal breast tissue. In all graphs, *, **, *** and **** symbols designate a significant difference of p<0.5, p<0.01, p<0.001 and p<0.0001, respectively.

**Supp. Fig 4.**
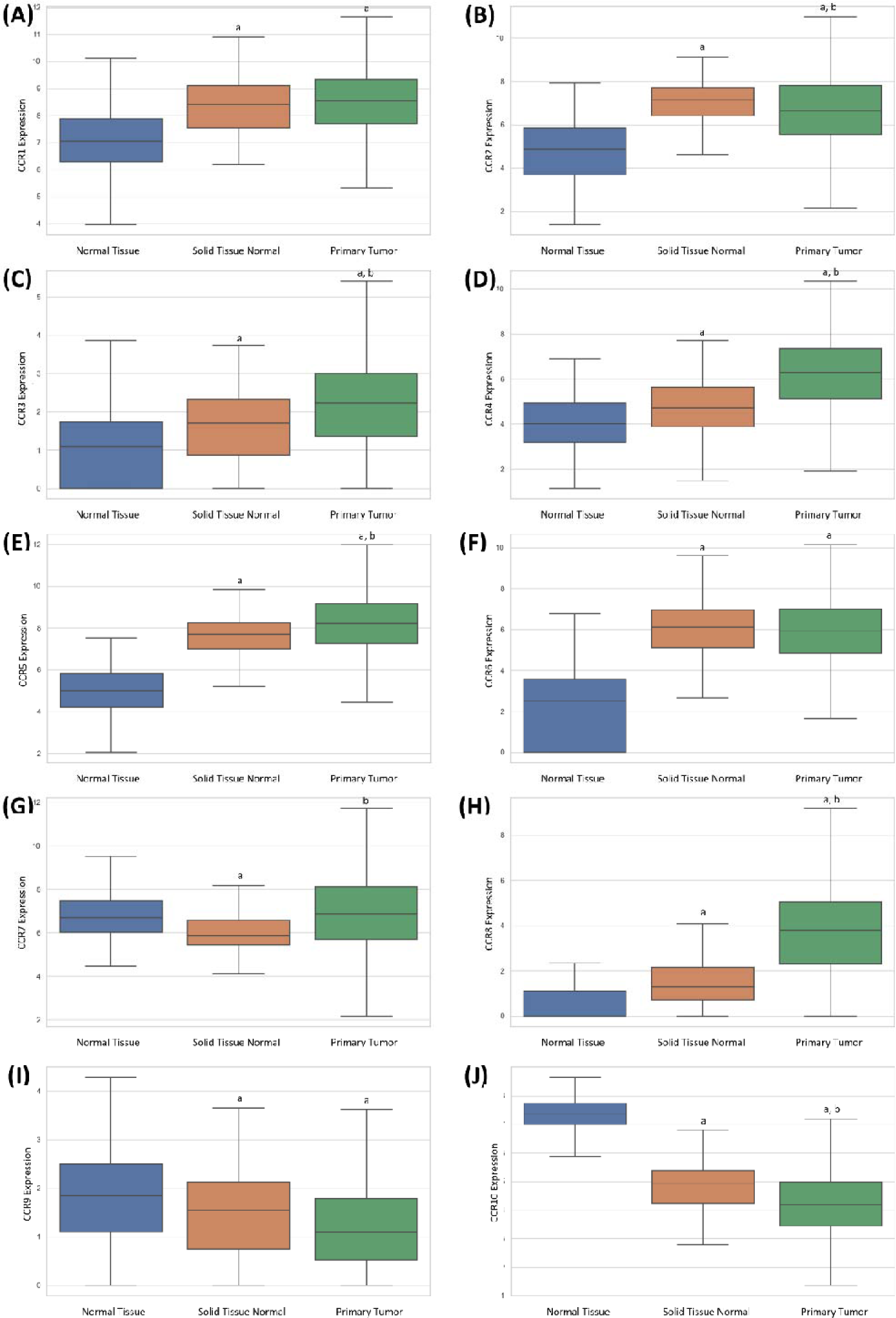
The expressions of CC Chemokine Receptors, CCRs 1-10 retrieved from Xena website and modified (n=178 for healthy normal tissue, n=113 for tumor-adjacent normal tissue, n=1092 for primary tumor). The unit of the expression values are log_2_(RSEM norm_count +1). **Note:** In all graphs, “a” and “b” symbols designate a significant difference of p<0.5 when compared to normal tissue and solid tissue normal, respectively based off of Tukey’s multiple comparison post hoc test.

**Supp. Fig 5.**
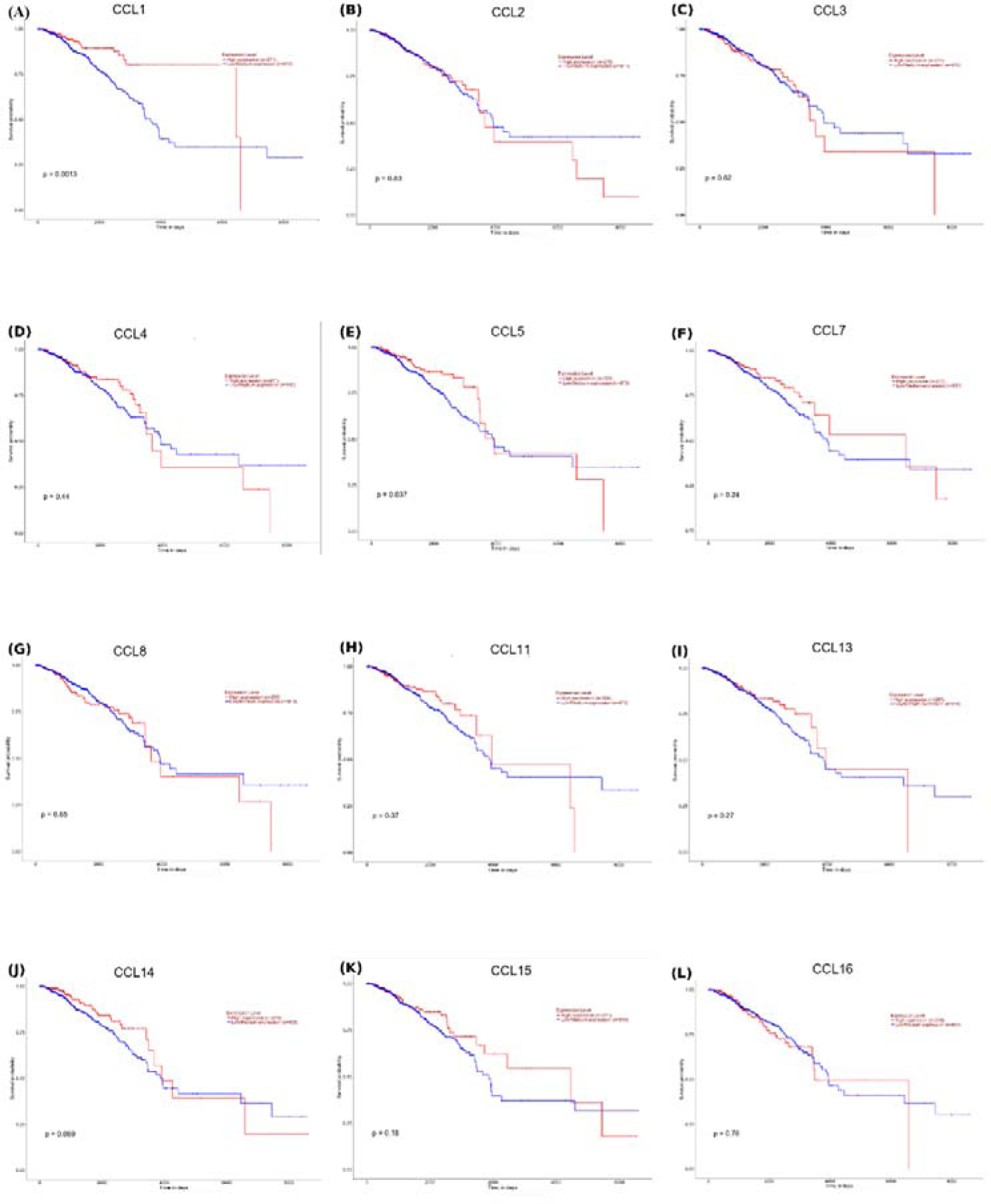
Patient Survival Chart of high (red color) versus low (blue color) expression levels of CCL1-5, CCL7-8, CCL11, and CCL13-16.

**Supp. Fig 6.**
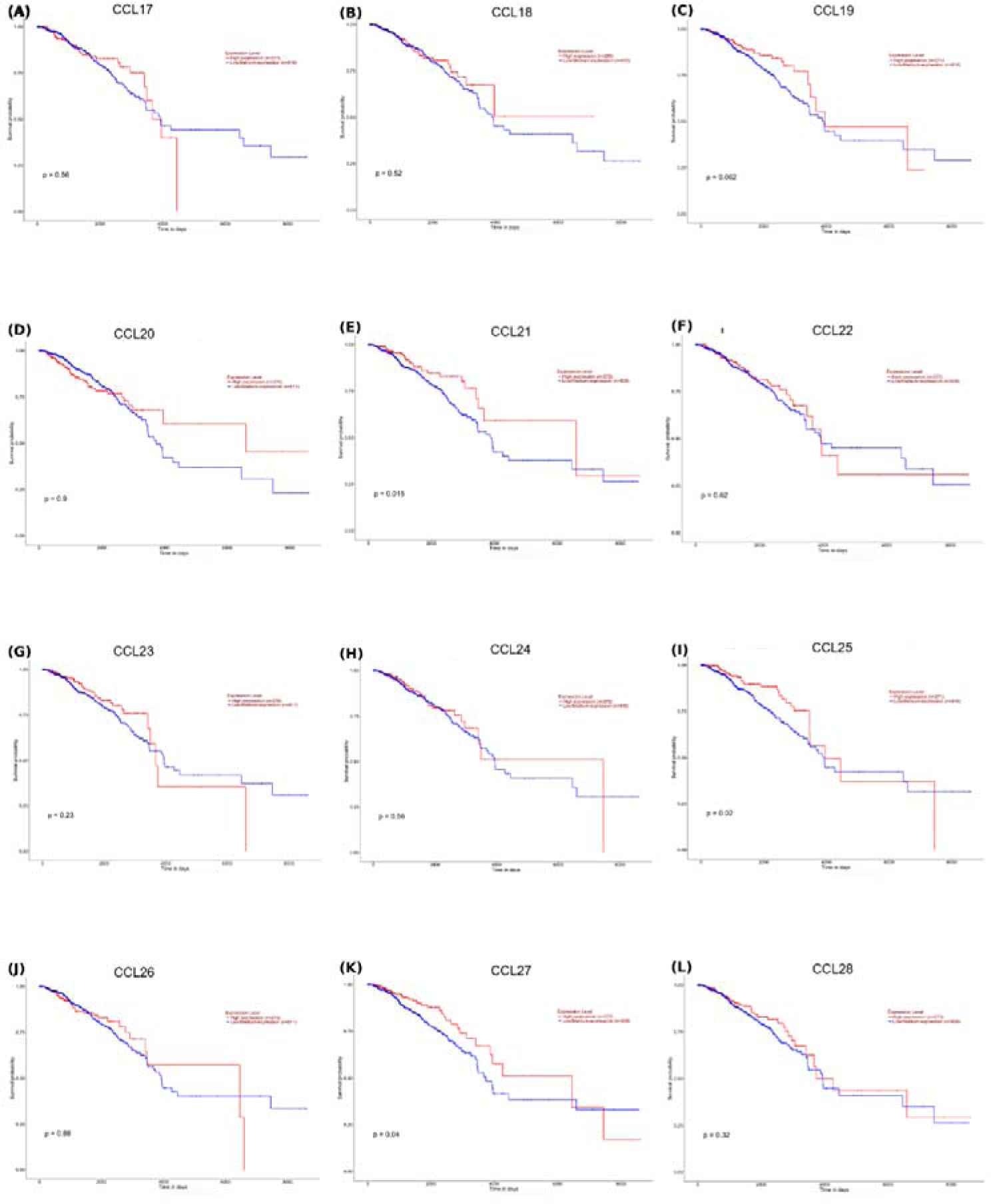
Patient Survival Chart of high (red color) versus low (blue color) expression levels of CCL17-25.

**Supp. Fig 7.**
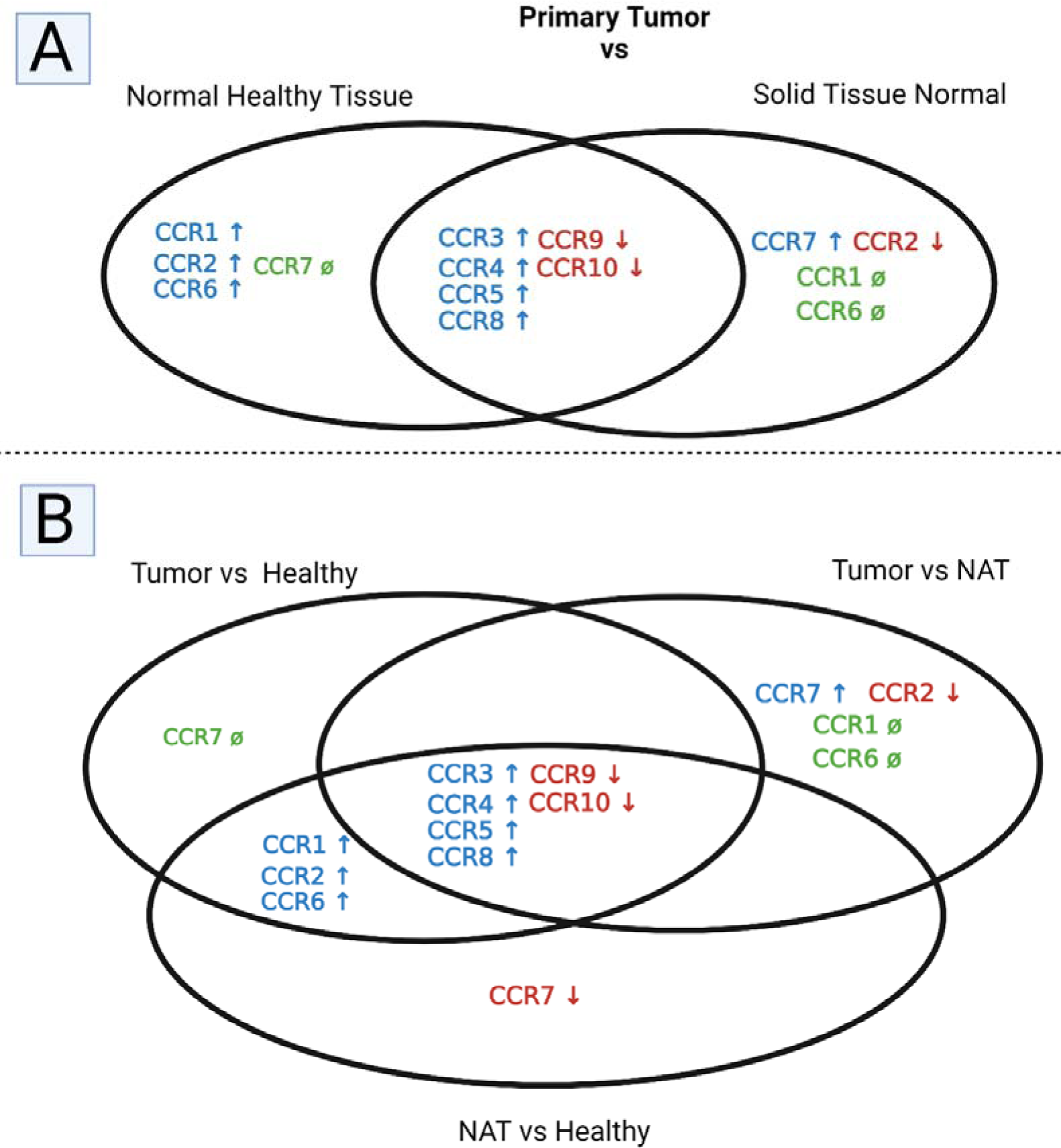
Venn Diagrams of the receptors of human CC-motif chemokines - CCRs 1-10: Comparing Expression Changes in Primary Tumor, Healthy Breast Tissue, and Normal Tumor Adjacent Tissue (NAT) (**A**) The Dual Venn Diagram of the overlapping similarities and distinctions in primary tumor tissue CCL expression changes over healthy breast tissue (left side) or NAT (right side). Green(ø), blue (↑) and red (↓) denote “no change”, “a significant increase” and “a significant decrease when compared to healthy (left side) or NAT (right side). (**B**) The triplet Venn Diagram of the overlapping similarities and distinctions in primary tumor tissue and NAT. In this diagram, primary tumor expression changes when compared to normal breast tissue (Tumor vs healthy) or tumor adjacent normal tissue (Tumor vs NAT) and NAT expression changes when compared to normal breast tissue (NAT vs Healthy) are shown. Green(ø), blue (↑) and red (↓) denote “no change”, “a significant increase” and “a significant decrease”, respectively.

**Supp. Fig 8.**
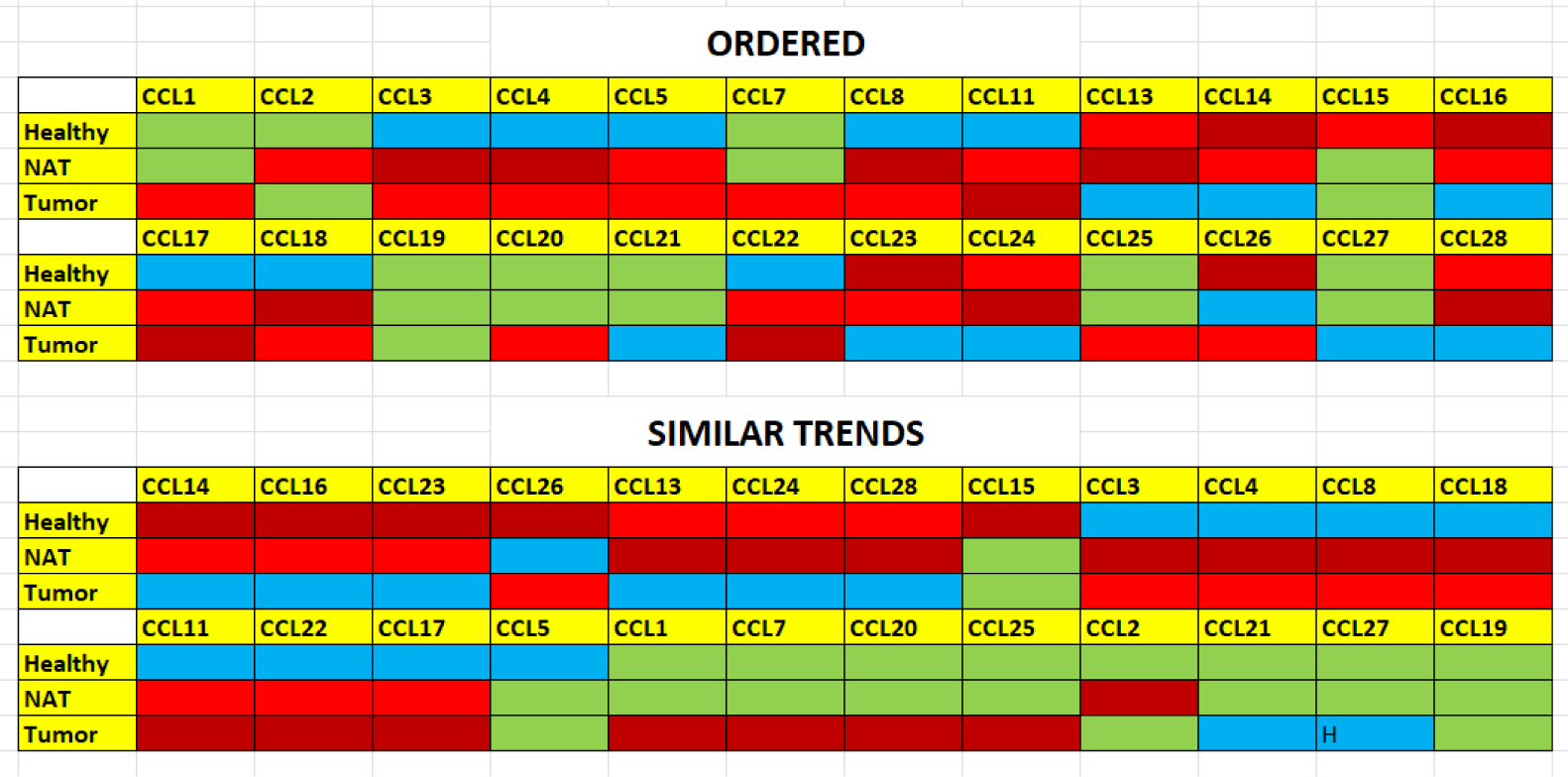
Colored illustration of expression values of different Human CC-type cytokines. Dark red denotes the highest expression, light red denotes the moderate expression, blue denotes lowest expression and green denotes similar expression between same colors.

## Notes

### Competing Interest Statement

The authors have declared no competing interest.

