## Supplementary figures and images for "CHEMOKINE SIGNALING IN BREAST CANCER: FOCUS ON CC-MOTIF CHEMOKINE LIGANDS"

### Supp. Fig 1

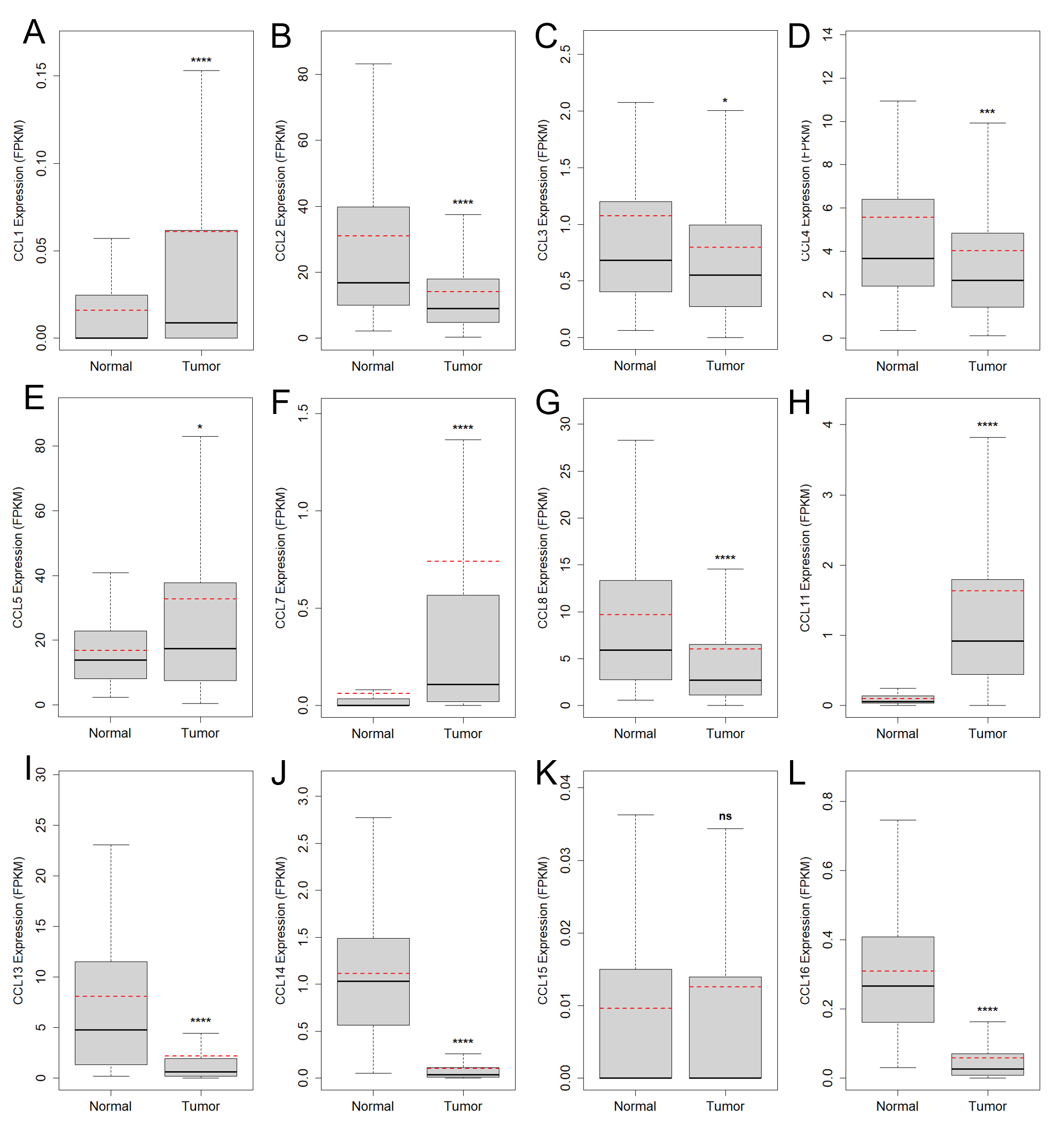

### Supp. Fig 2

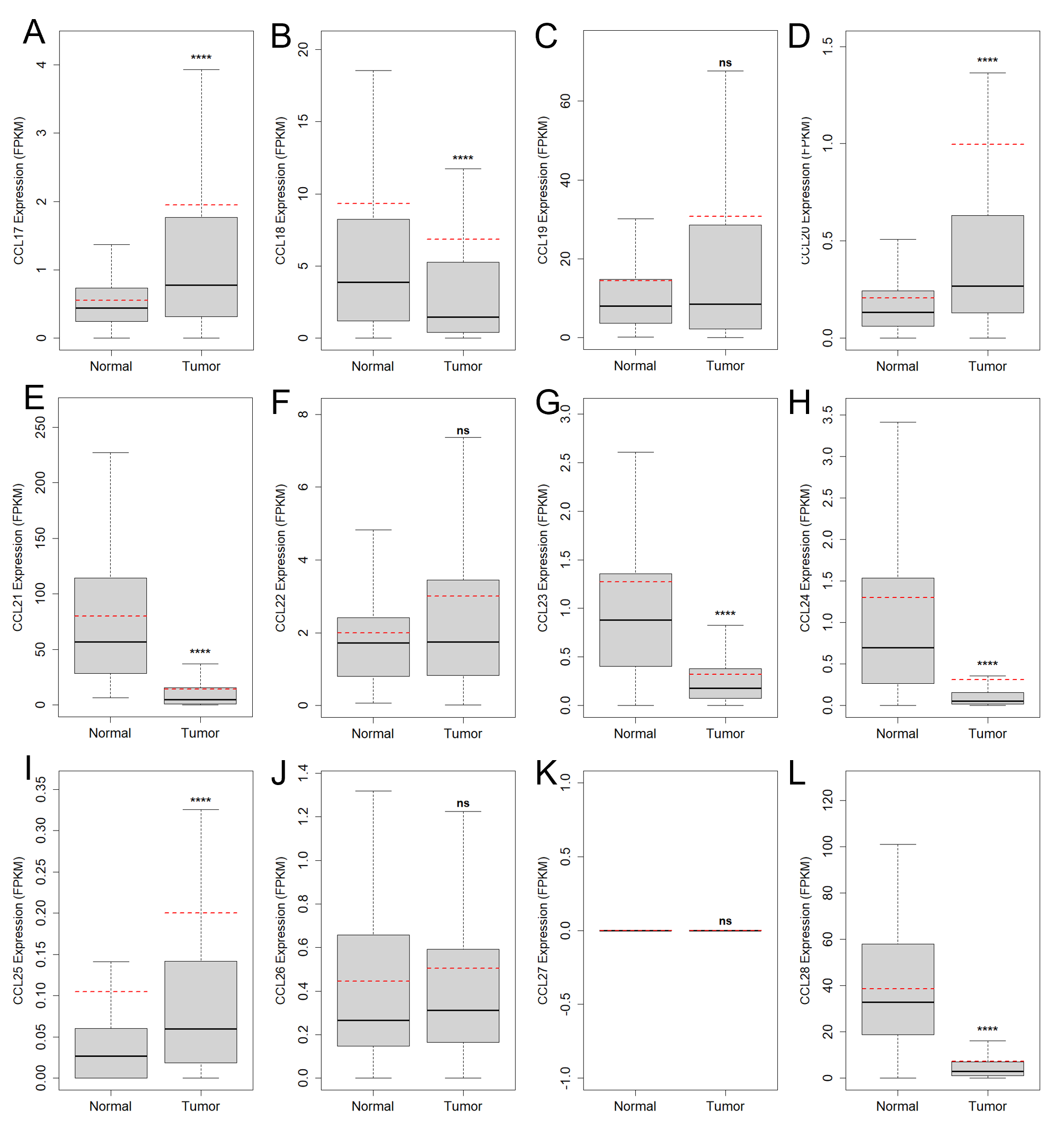

### Supp. Fig 3

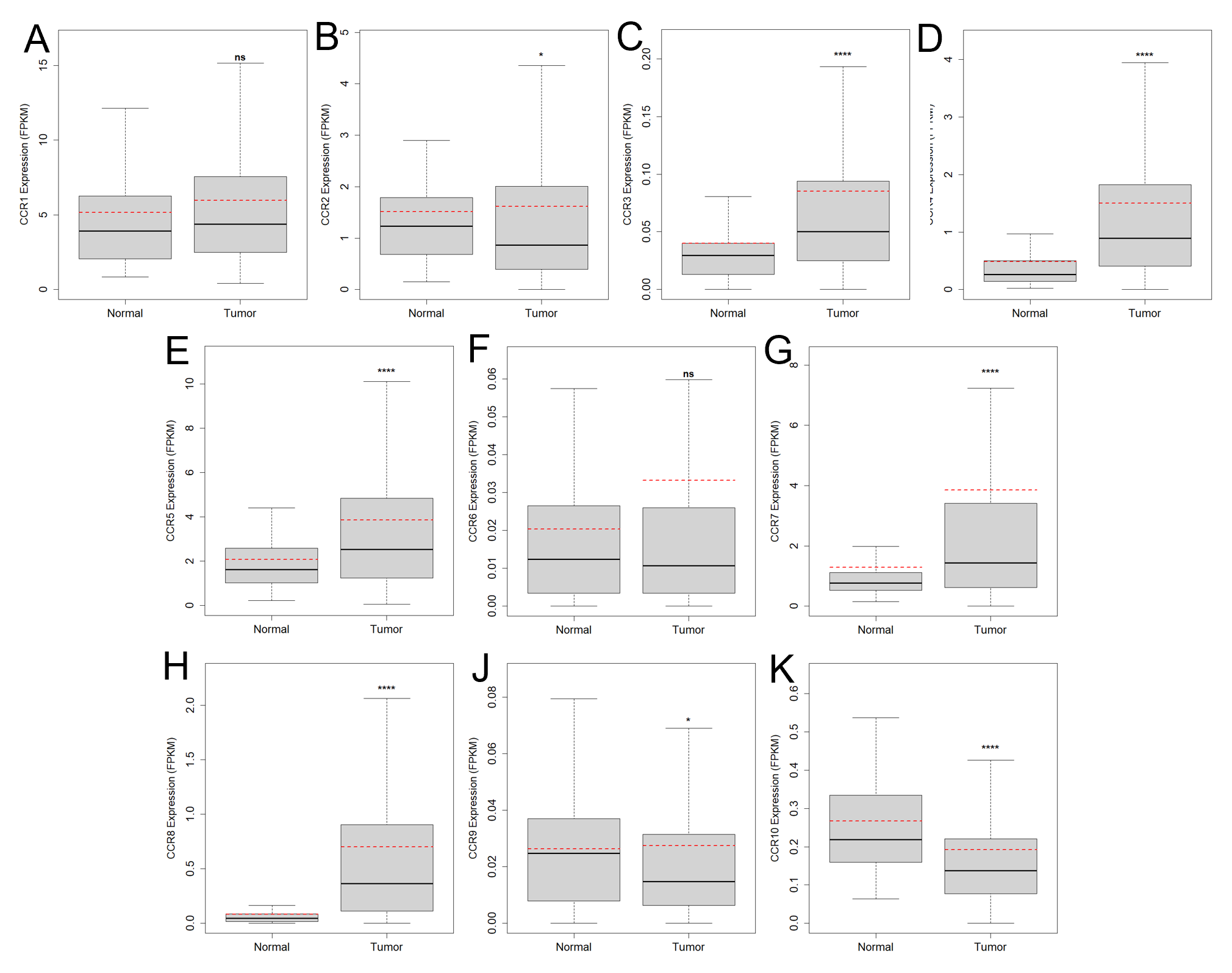

### Supp. Fig 4

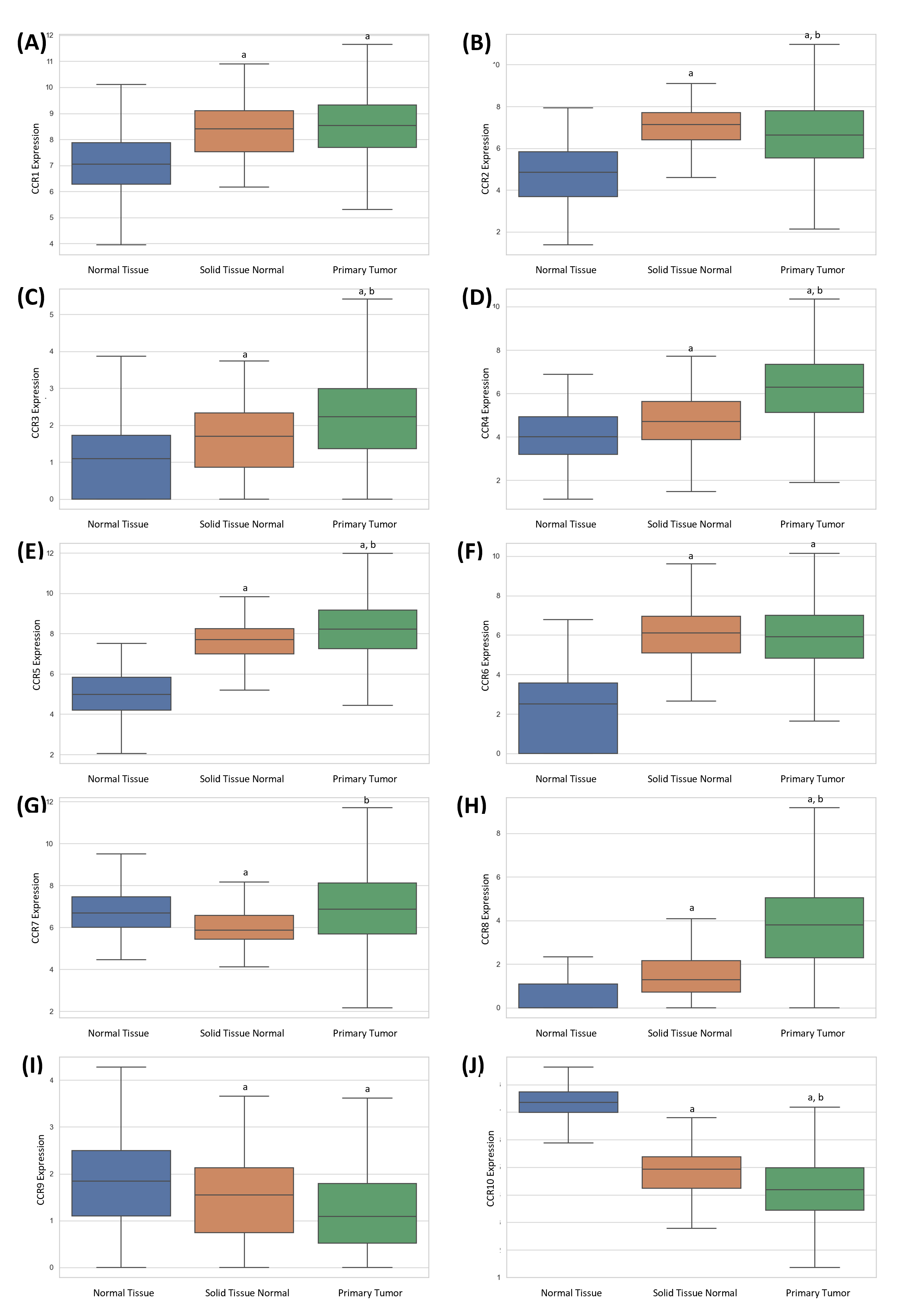

### Supp. Fig 5

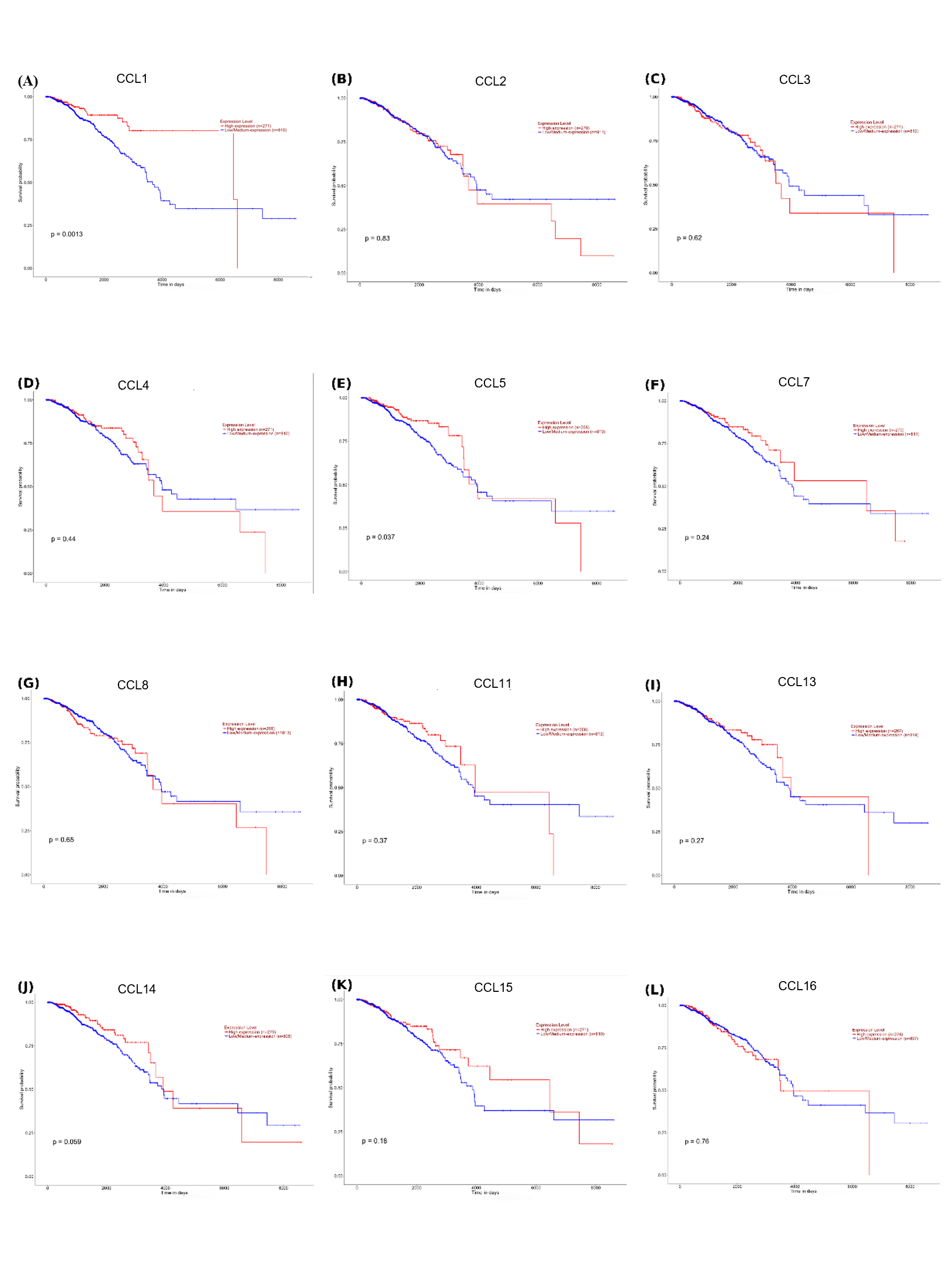

### Supp. Fig 6

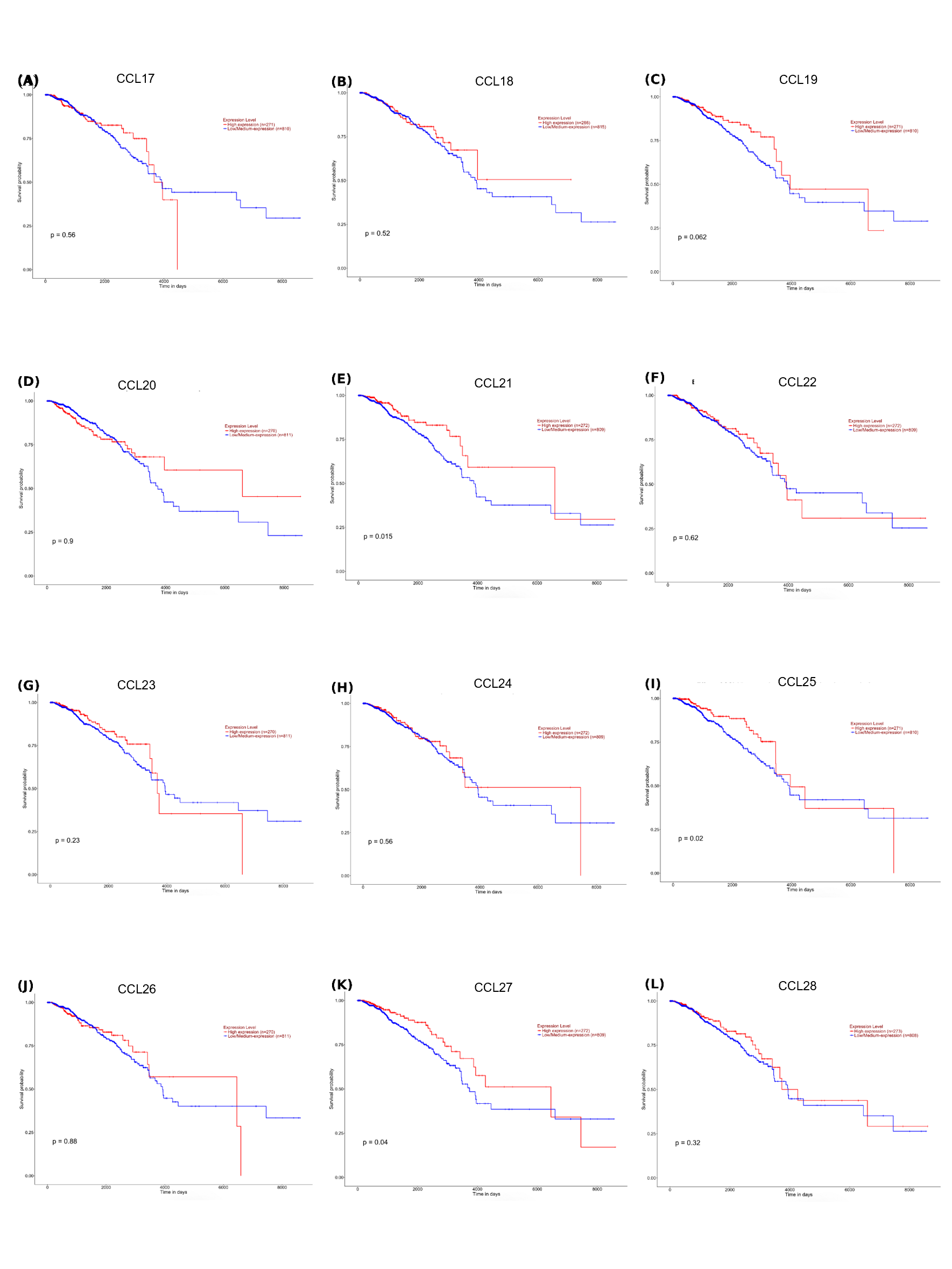

### Supp. Fig 7

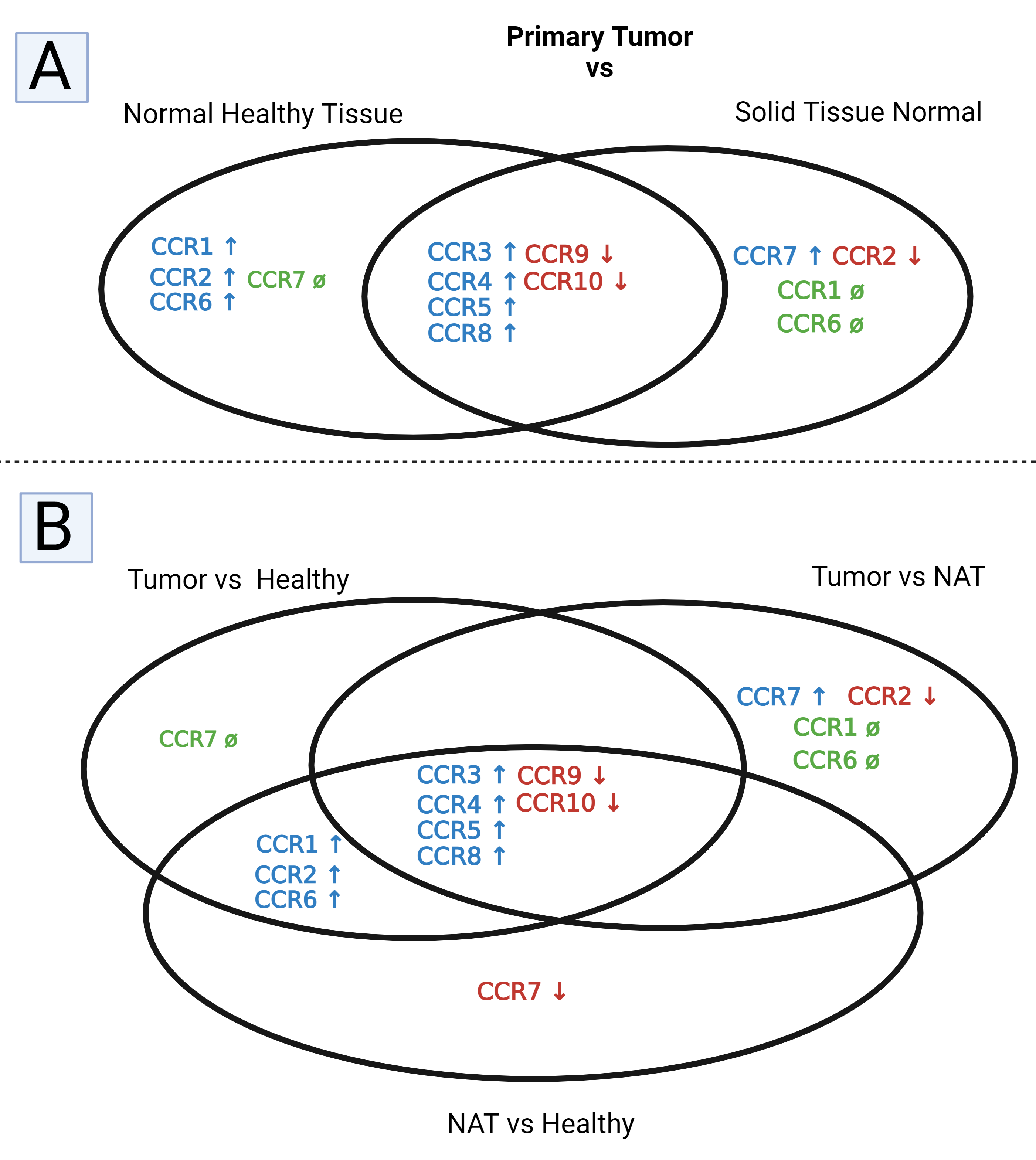

### Supp. Fig 8

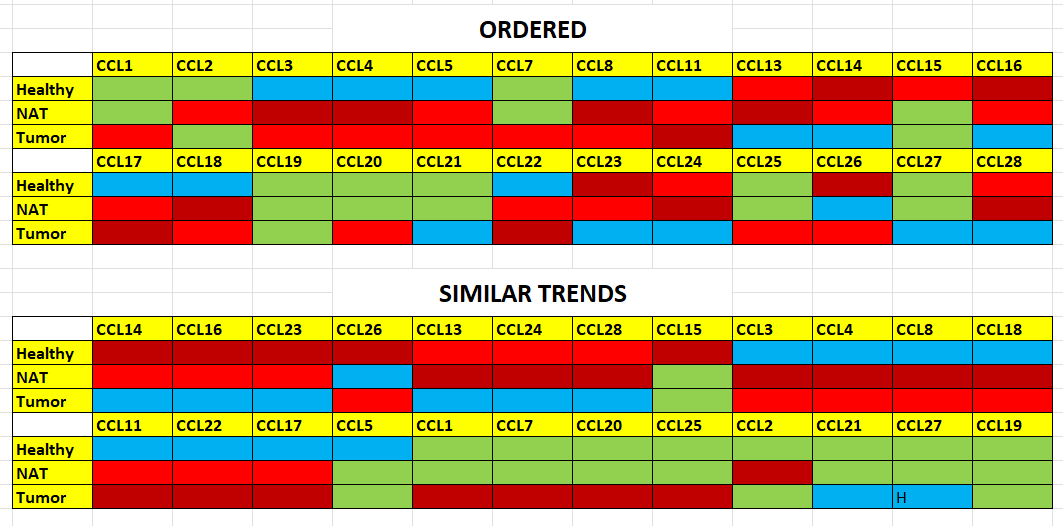
